# Conserved long-range base pairings are associated with pre-mRNA processing of human genes

**DOI:** 10.1101/2020.05.05.076927

**Authors:** Svetlana Kalmykova, Marina Kalinina, Stepan Denisov, Alexey Mironov, Dmitry Skvortsov, Roderic Guigó, Dmitri Pervouchine

## Abstract

The ability of nucleic acids to form double-stranded structures is essential for all living systems on Earth. While DNA employs it for genome replication, RNA molecules fold into complicated secondary and tertiary structures. Current knowledge on functional RNA structures in human protein-coding genes is focused on locally-occurring base pairs. However, chemical crosslinking and proximity ligation experiments have demonstrated that long-range RNA structures are highly abundant. Here, we present the most complete to-date catalog of conserved long-range RNA structures in the human transcriptome, which consists of 916,360 pairs of conserved complementary regions (PCCRs). PCCRs tend to occur within introns proximally to splice sites, suppress intervening exons, circumscribe circular RNAs, and exert an obstructive effect on cryptic and inactive splice sites. The double-stranded structure of PCCRs is supported by a significant decrease of icSHAPE nucleotide accessibility, high abundance of A-to-I RNA editing sites, and frequent occurrence of forked eCLIP peaks nearby. Introns with PCCRs show a distinct splicing pattern in response to RNA Pol II slowdown suggesting that splicing is widely affected by co-transcriptional RNA folding. Additionally, transcript starts and ends are strongly enriched in regions between complementary parts of PCCRs, leading to an intriguing hypothesis that RNA folding coupled with splicing could mediate co-transcriptional suppression of premature cleavage and polyadenylation events. PCCR detection procedure is highly sensitive with respect to *bona fide* validated RNA structures at the expense of having a high false positive rate, which cannot be reduced without loss of sensitivity. The catalog of PCCRs is visualized through a UCSC Genome Browser track hub.

## Introduction

Double-stranded structure is a key feature of nucleic acids that enables replicating the genomic information and underlies fundamental cellular processes (*1, 2*). Many RNAs adopt functional secondary structures, and mRNAs are no exception although their main role is to encode proteins (*3–6*). In eukaryotes, RNA structure affects gene expression through modulating all steps of pre-mRNA processing including splicing (*7*), cleavage and polyadenylation (*8*), and RNA editing (*9*). The loss of functional RNA structure has been increasingly reported as implicated in human disease (*10–13*).

To date, a few dozens of functional RNA structures have been characterized in the human genome (Tables 1 and S1). Many of them are formed between evolutionarily conserved regions that are located in introns of protein-coding genes and consist of base pairings spanning thousands of nucleotides. Computational identification of such distant base pairings by *de novo* RNA folding is not feasible due to a number of technical limitations (*14*). However, recent progress in high-throughput sequencing techniques enabled novel experimental strategies to determine RNA structure *in vivo* (*15–19*). In particular, photo-inducible RNA crosslinking and proximity ligation assays revealed that long-range base pairings are highly abundant in the human transcriptome (*20–24*). Currently, the applicability of these assays for large-scale profiling of RNA structure is still limited, and computational identification of long-range RNA structure remains a great challenge in RNA biology.

**Table 1:** Experimentally-validated (*bona fide*) RNA structures in human genes that satisfied PrePH search criteria. ∆*G* is the predicted free energy (kcal/mol); *l* is the average length of the two CCRs (nts); *d* is the spread (distance between complementary regions, nts); E-value is from R-scape after correction for multiple testing; *s*_1_, *s*_2_, and *s*_3_ are nucleotide conservation metrics (see Compensatory substitutions section).

| PCCR id | Gene | Exon | $\Delta G$ | $l$ | $d$ | E-value | $s_1$ | $s_2$ | $s_3$ | Refs |
| --- | --- | --- | --- | --- | --- | --- | --- | --- | --- | --- |
| 98636 | ENAH | exon 11a | -19.80 | 11 | 1728 | 0.4090 | 0.43 | 0.49 | 0.28 | (32) |
| 178925 | SF1 | exon 10 | -25.70 | 14 | 109 | 0.0002 | 0.45 | 0.49 | 0.64 | (28) |
| 739752 | DST | exons 46–52 | -25.00 | 15 | 9431 | 0.9998 | 0.39 | 0.56 | 0.45 | (29) |
| 883328 | DNM1 | exons 10a&b | -25.90 | 13 | 8864 | 0.2517 | 0.48 | 0.46 | 0.24 | (120) |
| 918502 | PLP1 | exon 3 | -15.80 | 20 | 638 | 0.3871 | 0.24 | 0.71 | 0.36 | (35) |
| 148879 | ATE1 | exon 7b | -33.2 | 17 | 51 | 0.6504 | 0.21 | 0.75 | 0.38 | (36) |
| 148881 | ATE1 | exon 7a | -35.9 | 13 | 952 | 0.5877 | 0.32 | 0.64 | 0.31 | (36) |

Comparative genomics provides a powerful alternative to *de novo* RNA folding by detecting signatures of evolutionary conservation (*25, 26*). Previous reports presented complex methodologies that implement simultaneous alignment and folding to detect RNA elements with divergent sequences that are nevertheless conserved at the secondary structure level (*27–30*). However, a substantial fraction (3.4%) of intronic nucleotides in the human genome are highly conserved at the sequence level, which raises a compelling question of whether their function may be related to RNA structure. This motivated us to revisit this problem with the “first-align-then-fold” approach (*14*), one which finds pairs of conserved complementary regions (PCCR) in pre-aligned evolutionarily conserved regions. In this study, we developed a method named PrePH to efficiently find long, nearly perfect complementary matches in a pair of input sequences, and applied it to all pairwise combinations of conserved intronic regions located at a certain distance limit from each other. Subsequently, we analyzed multiple sequence alignments that had a sufficient amount of variation to detect compensatory substitutions within PCCRs.

Multiple lines of evidence indicate that a large proportion of PCCRs indeed have a double-stranded structure, e.g., significant decrease of icSHAPE nucleotide accessibility, high abundance of A-to-I RNA editing sites, significant overlap with long-range RNA contacts identified by proximity ligation assays, and frequent co-occurrence of the so-called forked eCLIP peaks (see below). They also have other characteristic features such as occurrence within introns proximally to splice sites, avoidance of branch points, and obstructive effect on cryptic and inactive splice sites. At the same time, the method has a substantial false positive rate which, as we show, cannot be reduced without losing sensitivity to *bona fide* validated RNA structures (Table 1). The catalog of PCCRs is provided as a reference dataset that is conveniently visualized through a UCSC Genome Browser track hub (*31*). We additionally provide a transcriptome-wide characterization of RNA bridges (*32*) and exon loop-outs (*33*), two particular mechanisms of alternative splicing regulation by long-range RNA structures.

## Results

### Conserved Complementary Regions

To identify conserved complementary regions (CCR), we considered nucleotide sequences of human protein-coding genes excluding all constitutive and alternative exons, repeats, non-coding genes residing in introns, and other regions with selective constraints that may not be related to base pairings (Figure 1A). The remaining intronic regions were extended by 10 nts to within flanking exons to allow for base pairings that overlap splice sites. The resulting set of 236,332 intronic regions was intersected with the `phastConsElements` track of the UCSC Genome Browser (*34*), which defines genomic intervals that are highly conserved between 100 vertebrates (major species *P. troglodytes, M. musculus, S. scrofa, G. gallus, X. tropicalis, D. rerio*; the shortest and the longest phylogenetic distances 0.01 and 2.40, respectively). This resulted in a set of 1,931,116 short fragments with the median length 17 nts, which will be referred to as conserved intronic regions (CIR).

Pairs of conserved complementary regions (PCCRs) were identified in all pairwise combinations of CIR that are located not more than *L* nts apart from each other and belong to the same gene using PrePH, a *k*-mer-based method (Figure 1B) that efficiently predicts long, nearly perfect stretches of complementary nucleotides in a pair of input sequences (see Methods). A search for at least 10-nt-long sequences with the hybridization free energy ∆*G* ≤ −15 kcal/mol, minimum helix length *k* ≥ 5, and distance limit *L* ≤ 10, 000 yielded 916,360 PCCRs, on average 75 PCCR per gene, with 95% of genes having not more than 295 PCCRs (Supplementary Data Files 1 and 2). The median free energy of hybridization (∆*G*) and the median length of CCRs were −17.2 kcal/mol and 13 nts, respectively, with the frequency distribution decaying towards longer and more stable structures (Figure 1C and S1A). As expected, longer structures had larger absolute values of ∆*G*; however, the energy and the length of a PCCR are not directly proportional and the relationship between them depends on the GC content (Figures S1B and S1C). In what follows, PCCRs are classified into four energy groups, group I from −15 to −20 kcal/mol, group II from −20 to −25 kcal/mol, group III from −25 to −30 kcal/mol, and group IV below −30 kcal/mol, which are represented throughout the paper by a uniform color scheme shown in Figure 1C.

**Figure 1:**
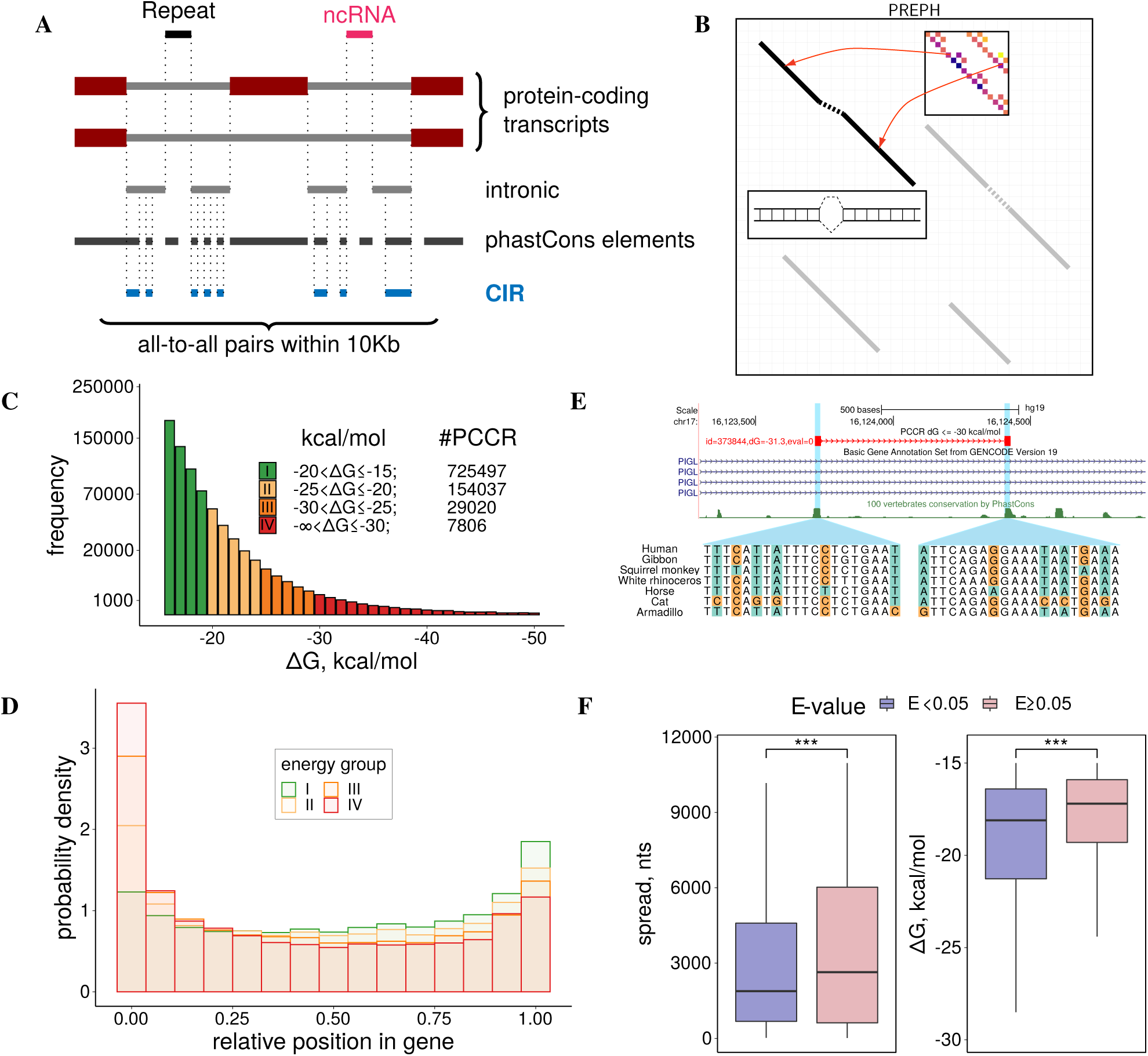
Pairs of Conserved Complementary Regions (PCCR). **(A)** PCCRs are identified in conserved intronic regions (CIR) that are less than 10,000 nt apart from each other. **(B)** PrePH computes the dynamic programming matrix based on the pre-computed helix energies for all *k*-mers (inset) and energies of short internal loops and bulges (see Supplementary Methods for details). **(C)** The distribution of PCCR energies consists of four energy groups: group I (−20 < ∆*G* ≤ −15 kcal/mol), group II (−25 < ∆*G* ≤ −20 kcal/mol), group III (−30 < ∆*G* ≤ −25 kcal/mol), and group IV (∆*G* ≤ −30 kcal/mol). **(D)** The distribution of *p*, relative position of a PCCR in the gene. **(E)** Multiple independent compensatory substitutions support long-range RNA structure in the phosphatidylinositol glycan anchor biosynthesis class L (PIGL) gene. **(F)** PCCRs with significant nucleotide covariations (E-value < 0.05) are on average less spread and more stable than other PCCRs.

To assess the sensitivity of the method with respect to known RNA structures, we collected evidence of functional base pairings in human genes from the literature. All *bona fide* structures that satisfied the search criteria were successfully found (Table 1, Figure S2), while others either didn’t pass the free energy cutoff, were shorter than 10 nts, located outside of CIR, or didn’t belong to protein-coding genes (Table S1). Notably, the long-distance intronic interaction that regulates *PLP1/DM20* splicing (*35*) and the RNA bridge in *ENAH* (*32*) were both assigned to group I (∆*G* = −15.8 and −19.4 kcal/mol, respectively) indicating that less stable structures are not less functional or less interesting than the others. For the purpose of presentation here, we chose the distance limit *L* = 10, 000, which by the order of magnitude corresponds to long-range RNA structures listed in Table 1. However, our recent study of the human *ATE1* gene demonstrated that functional RNA structures may spread over much longer distances (*36*). We therefore additionally explored how the number of PCCRs changes with increasing the distance limit and found that it grows approximately fourfold with increasing *L* up to 100,000 nts (Figure S3A).

The frequency distribution of the distances between two CCRs in a pair, here referred to as *spread*, peaks at short distances, decaying towards a non-zero baseline (Figure S3B). This baseline originates from the distribution of pairwise distances between CIR that were passed to PrePH as an input, which decays towards the same baseline. We next asked whether PCCRs were distributed uniformly along the gene, or they tend to accumulate in certain gene parts. To quantify the position of a PCCR within a gene, we introduced *p*, a measure of relative position, which changes from 0% for the regions located in the very 5’-end of the gene to 100% for the regions located in the very 3’-end. The metric *p* can be computed for a PCCR as a whole to represent its relative position, or for each of its CCR separately. The location of PCCR as a whole was not uniform, with two pronounced modes at the 5’- and 3’-end (Figure 1D). This enrichment was also prominent in the distribution of single CCRs (Figure S3C), which could be due to stronger evolutionary constraints on the nucleotide sequences at gene ends. Indeed, we found that CCRs exhibit a higher degree of evolutionary conservation than their adjacent regions within CIR (the difference of the average phastCons scores, MW test, p-value < 2.2∗10^−16^) and thus tend to occur in more constrained regions. Gene ontology terms associated with genes that have PCCRs were enriched with terms related to morphogenesis and development of the central nervous system as compared to genes of the same length, but without PCCRs (Figure S4).

### Compensatory substitutions

Compensatory mutations, i.e., pairs of nucleotide substitutions that individually disrupt base pairings, but restore them when introduced in combination, play a central role in the evolution of RNA structure [7568070, 12590655, 15502829]. To analyze compensatory mutations in PCCRs, we applied the R-scape program (*37*), which scores independent occurrence of complementary substitutions on different branches of the phylogenetic tree, to pairs of multiple sequence alignments (MSA) that were cut out from the MSA of 100 vertebrates (*34*) by PCCRs (see Methods). The deviation from the null hypothesis that pairwise covariations in a PCCR are not due to conservation of RNA structure was estimated as a product of E-values reported by R-scape for all base pairs within the PCCR. These products were adjusted using Benjamini-Hochberg correction.

Out of 916,360 PCCR, for which this computation was possible, only 909,146 had a sufficient number of substitutions to estimate the E-value, and only 3,204 of them had E-value below 5% (Figure S5). The PCCRs with E-value < 0.05 were on average more stable and less spread than PCCRs with E-value ≥ 0.05 (Figure 1F). In some cases, structural alignments were strongly supported by covariations, i.e., a highly stable PCCR (∆*G* = −31 kcal/mol) spanning 700 nts in the first intron of *PIGL* gene (Figure 1E) and PCCRs in the *QRICH2* and *MRPL42* genes (Figure S6). However, E-values of *bona fide* RNA structures from Table 1 ranged from 0.0002 for the PCCR responsible for splicing of the intron between exons 9 and 10 in *SF1* to 0.9998 for the PCCR associated with exon 46–52 skipping in *DST*, suggesting that the amount of variation in the nucleotide sequences of PCCRs is generally not sufficient to estimate their statistical significance through compensatory substitutions.

A remarkable feature of standalone RNA regulatory elements is their high level of conservation, as opposed to that of their flanking sequences (*14*). We introduced three additional metrics to capture the nucleotide conservation rate within CCR as compared to the background: *s*_1_ is the difference between the average phastCons scores within CCR and within 300 nt flanking regions (the larger, the more significant); *s*_2_ is the average phastCons score within CCR and 300 nt around it (the smaller, the more significant); *s*_3_ is the length of a CCR relative to the length of its parent CIR (the larger, the more significant). However, neither of the three metrics reached extreme values for the base pairings listed in Table 1 (Figure S7), which indicates that functional PCCRs are not necessarily located in isolated conserved regions and may well occur in a relatively conserved background.

### Support by high-throughput structural assays

PCCRs represent regions in the pre-mRNA that have an increased capacity to base-pair. The propensity of individual nucleotides to base-pair was assessed at the transcriptome-wide level by measuring the nucleotide flexibility score with icSHAPE method (*38*). We compared the average icSHAPE reactivity within CCR with that in a control set of intervals located nearby (Figure 2A). Indeed, the average icSHAPE reactivity of nucleotides within CCR was significantly lower as compared to the control (Wilcoxon test, P < 10^−60^), and the difference increased by the absolute value with increasing the structure free energy 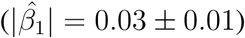. However, the icSHAPE reactivity scores were available only for 4,551 PCCRs representing 0.5% of the full set. We therefore sought for PCCR support in other experimental datasets.

**Figure 2:**
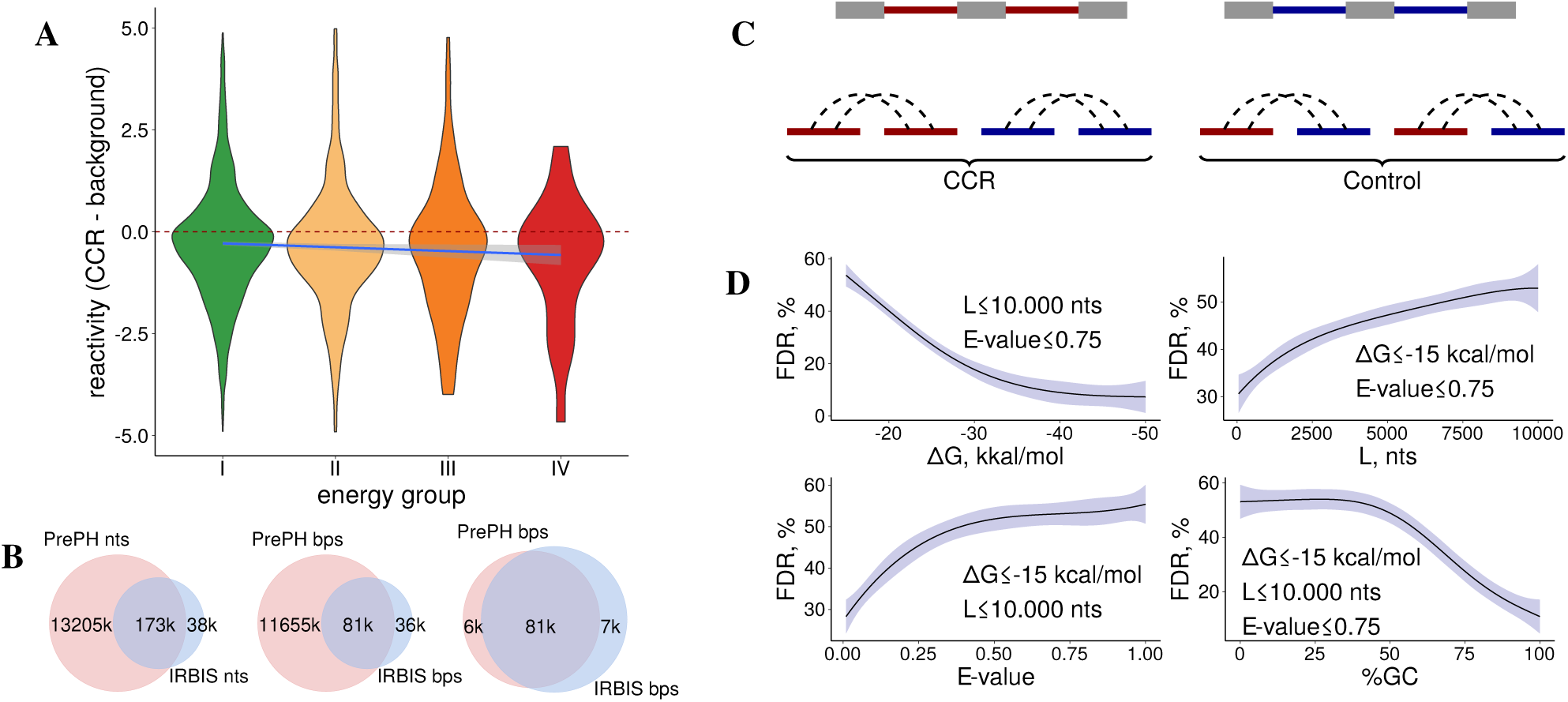
Validation and False Discovery Rate (FDR). **(A)** The difference between icSHAPE reactivity of nucleotides within CCR and the average reactivity of nearby nucleotides in energy groups I–IV (color code as in Figure 1C). The linear model ∆*reactivity* = *β*_0_ + *β*_1_∆*Ggroup* is represented by the slanted line; *β̂*_1_ = −0.03 ± 0.01. **(B)** Venn diagram for the number of common nucleotides (left), number of common base pairs (middle), and the number of common base pairs among common nucleotides (right) for the predictions of PrePH and IRBIS. **(C)** Estimation of the false positive rate (FDR) by rewiring, i.e., creating a control set that consists of chimeric non-cognate sequences sampled from different genes. **(D)** FDR as a function of energy cutoff ∆*G* (top left), maximum distance between CIR (top right), E-value (bottom left), and GC content (bottom right). Shaded areas represent 95% confidence intervals in *n* = 16 randomizations.

Chemical RNA structure probing can reveal which bases are single- or double-stranded, but it cannot determine which nucleotides form base pairs (*15–19*). We therefore validated PCCRs against the long-range RNA-RNA interactions that were assessed experimentally by PARIS (*22*) (*n* = 15, 036), LIGR-seq (*23*) (*n* = 551, 926), and RIC-seq (*24*) (*n* = 501, 144). Towards this end, we considered the interacting pairs in the experimental data that were located intramolecularly within CIR not more than 10,000 nt apart from each other, and restricted the set of PCCRs to the underlying CIRs that overlap the experimental dataset (see Methods). The precision (*P*) and recall (*R*) metrics were defined as the proportion of PCCRs supported by the experimental method and the proportion of experimental interactions supported by PCCRs, respectively. Additionally, we computed *π*, the conditional probability of predicting the interacting CCR partner correctly given that another CCRs in a pair has been predicted correctly (Table 2). The best agreement was with respect to the RIC-seq dataset, with precision increasing up to 92% at the expense of decreasing recall when structure stability increased. The *π* metric confirmed that PCCRs tend to correctly identify the interacting partner given that one of the CCRs has been predicted correctly. Additionally, we found that free energies of PCCRs supported by RIC-seq and PARIS were significantly lower than those of PCCRs without experimental support (MW test, ∆∆*G* ≃ 1.2 kcal/mol, P-value < 10^−19^). However, the breadth of these findings is limited by small sizes of the true positive sets (*n* = 1, 903 for RIC-seq, *n* = 777 for LIGR-seq, and *n* = 969 for PARIS), because structural assays sparsely cover the transcriptome at cell-line-specific conditions and focus on intermolecular interactions.

**Table 2:** Precision and recall at different free energy cutoffs (∆*G*). The precision (*P*) and recall (*R*) are the proportion of PCCRs supported by the experimental method and the proportion of experimental interactions supported by PCCRs, respectively. *π* is the conditional probability of predicting the interacting CCR partner correctly given that another CCRs in a pair has been predicted correctly. The column ‘PrePH’ shows the number of PCCRs that satisfy the criteria for comparison. The number of structures in each experimental method is denoted by *n*.

|  | **RIC-seq, $n = 1,804$** |  |  |  | **LIGR-seq, $n = 586$** |  |  |  | **PARIS, $n = 907$** |  |  |  |
| --- | --- | --- | --- | --- | --- | --- | --- | --- | --- | --- | --- | --- |
| $\Delta G$ | PrePH | $P, \%$ | $R, \%$ | $\pi, \%$ | PrePH | $P, \%$ | $R, \%$ | $\pi, \%$ | PrePH | $P, \%$ | $R, \%$ | $\pi, \%$ |
| -15 | 1611 | 42 | 53 | 43 | 362 | 54 | 49 | 56 | 5901 | 50 | 60 | 51 |
| -20 | 364 | 66 | 26 | 67 | 88 | 57 | 17 | 58 | 1725 | 60 | 33 | 60 |
| -25 | 84 | 86 | 9 | 86 | 21 | 48 | 3 | 48 | 545 | 65 | 14 | 65 |
| -30 | 23 | 91 | 3 | 91 | 5 | 40 | 1 | 40 | 160 | 72 | 6 | 73 |

Finally, we explored how the set of predicted PCCRs relates to a similar list that was reported previously by IRBIS (*29*). Unlike PrePH, IRBIS follows the “first-fold-then-align” strategy to simultaneously detect conserved complementarity and sequence homology, but at stricter conditions. Overall, the predictions of the two programs had a large intersection relative to IRBIS predictions indicating that the current method generally outputs a superset of IRBIS predictions both in terms of the number of nucleotides and the number of base pairs (Figure 2B). The free energies of PCCRs supported by IRBIS were significantly lower than those of other PCCRs (MW test, ∆∆*G* ≃ 2.7 kcal/mol, P-value < 10^−30^) reflecting the fact that, unlike IRBIS, PrePH allows for short internal loops and bulges. Nevertheless, a small fraction of IRBIS predictions are missing from the list of PCCRs presented here, which relies on the conserved regions from `phastConsElements` track of the UCSC Genome Browser (*34*). Without this limitation, however, the current approach would be impractical from the computational standpoint, and it was our intention to limit the method to conserved regions at the expense of losing some structures that are misaligned by structure-agnostic phylogenetic analysis.

### False Discovery Rate

One way to estimate the rate of false positive predictions is to apply the same pipeline that was used to identify PCCRs to a control set of sequences that should not base-pair. In the previous work, it was described as the so-called “re-wiring” approach (*29*). Here, we used the same strategy by running the pipeline on chimeric sets of sequences that were sampled randomly from different genes while controlling for nucleotide composition and length, which confound this comparison (Figure 2C). The false discovery rate (FDR), defined as the number of predictions in the control set as a fraction of the total number of predictions, depends on PCCR energy, spread, GC content, and E-value, ranging from 10% in the most strict to over 50% in the most relaxed conditions (Figure 2D). As expected, FDR drops with increasing PCCR energy and GC content and with decreasing PCCR spread and E-value.

Many PCCRs listed in Table 1 belong to group I indicating that folding energy alone cannot be used as a threshold to control FDR. To check whether FDR can be improved by simultaneous application of several filters, we applied the most stringent thresholds that preserve PCCRs listed in Table 1 (∆*G* ≤ −15.8, E-value < 0.998, *GC* ≥ 33.3%, *s*_1_ ≥ 0.24, *s*_2_ ≤ 0.71, and *s*_3_ ≥ 0.24). The resulting FDR figure of 47% indicates that FDR cannot be improved without loss of sensitivity with respect to *bona fide* structures. On the one hand, it implies that more than a half of the group I predictions could be false positives. On the other hand, this is a very pessimistic estimate since the re-wiring method is known to greatly overestimate the FDR (*29*). We therefore interpret 47% as an excessively conservative FDR estimate and proceed to the statistical characterization of the full PCCR set with the mindset that at least a half of the predictions are true positives.

### Splicing

Previous reports indicate that long-range base pairings are positioned non-randomly with respect to splicing signals (*27–29*). To elaborate on this, we used the classification shown in Figure 3A. If a PCCR overlaps an intron, it can be located either entirely within the intron (inside), or the intron can be located entirely within PCCR (outside), or the two intervals intersect (crossing). The tendency of RNA structure to prefer one of these categories is measured by the *enrichment* metric, defined as the number of PCCRs in the given category relative to the number of PCCR-like intervals in it, computed for a certain control set (each PCCR may be counted more than once if it is located inside one intron and crosses another). In the first control set, referred to as *random shift*, each PCCR was shifted randomly within its gene. The resulting pseudo-PCCR had the same spread and belonged to the same gene as the original PCCR. In the second control set, referred to as *random gene*, for each gene and a PCCR in it, we created a pseudo-PCCR at the same relative position as the original PCCR, but in a randomly chosen gene of the same length. The resulting pseudo-PCCR had the same spread and the same relative position as the original PCCR, but belonged to a different gene. Repeated sampling of these sets allowed estimation of statistical significance.

**Figure 3:**
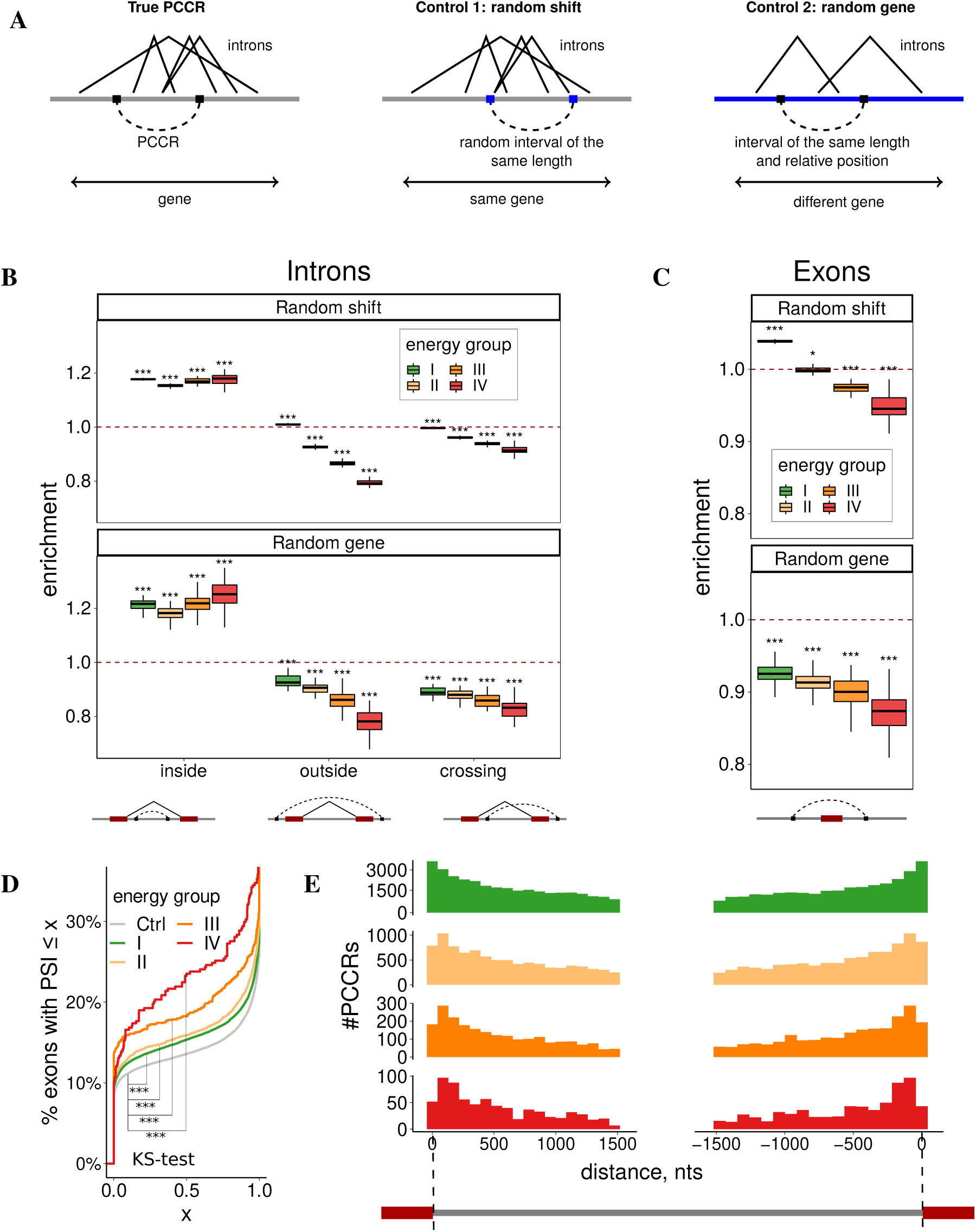
Splicing. **(A)** Control procedures. In the random shift control, a PCCR is shifted within the gene. In the random gene control, a pseudo-PCCR is created in the same relative position of a different gene chosen at random. The numbers of PCCRs inside, outside, and crossing the reference set of intervals (e.g., introns) are counted. **(B)** PCCRs are enriched inside introns and depleted in outside and crossing configurations. Boxplots reflect *n* = 40 randomizations. **(C)** PCCRs looping-out exons are depleted. **(D)** The cumulative distribution of the average exon inclusion rate (Ψ) in HepG2 cell line for exons looped-out by PCCRs of the four energy groups vs. exons not looped-out by PCCRs (Ctrl). **(E)** The distribution of distances from intronic PCCRs to intron ends (bin size 75 nts). Group I PCCRs are enriched, while group IV PCCRs are depleted in 75-nt windows immediately adjacent to splice sites.

In comparison with the set of annotated introns by random shifts, PCCRs showed a pronounced enrichment in the inside category and depletion in the outside and crossing categories, with the magnitude of depletion increasing for more stable structures (Figure 3B, top). To rule out a possibility that preferential PCCR positioning within introns originates from uneven distribution of longer introns along the gene, with 5’-introns being on average longer (*27, 28*), we repeated the same analysis using random gene control, and the enrichment and depletion remained significant (Figure 3B, bottom). A similar comparison with the set of annotated exons revealed a substantial depletion of PCCRs that loop-out exons, which also became stronger as PCCR stability increased (Figure 3C). The average inclusion rate of exons that were looped out by PCCR was lower than that of exons that are not surrounded by PCCR, with the magnitude of the difference increasing for more stable structures (Figure 3D). These results reconfirm that PCCRs generally tend to avoid placing exons in a loop, which promotes exon skipping.

The frequency of intronic PCCRs decays with increasing the distance to intron ends in all four energy groups partially reflecting the decrease of sequence conservation (Figure 3E). However, weaker structures occur more frequently in 75-nt windows adjacent to splice sites, while stronger structures tend to occupy more distant positions. The ends of PCCRs tend to be closer to exon boundaries than would be expected by chance from random shifts (Figure S8). Contrary to what was reported earlier for *D. melanogaster*, there is no substantial depletion of CCR in polypyrimidine tracts (PPT). This indicates a large regulatory potential for splicing since a large fraction of CCR that overlap PPT (43.2%) also block the acceptor splice site. In addition, there were on average 20% less overlaps of CCR with intronic branch points (*39*) than would be expected from random shift control, i.e., CCRs tend not to overlap intronic branch points.

To test whether PCCRs interfere with splicing signals, we identified intronic motifs with high similarity to donor and acceptor site consensus sequences (cryptic splice sites) and analyzed the expression of splice sites in a large compendium of RNA-seq samples from the Genotype Tissue Expression Project (GTEx) (*40*). CCRs overlapping highly expressed (active) splice sites were depleted, while CCRs overlapping splice sites with low read support (inactive) were enriched (Figure 4A). CCRs overlapping non-expressed cryptic splice sites were also enriched with the exception of highly stable structures, which are likely devoid of cryptic splice sites due to high GC content. We also found that the largest enrichment of CCRs overlapping candidate cryptic splice sites was among PCCR with small spread (MW test, p-value = 0.004), in agreement with the previous findings that cryptic splice sites tend to be suppressed by local RNA secondary structure (*41, 42*).

**Figure 4:**
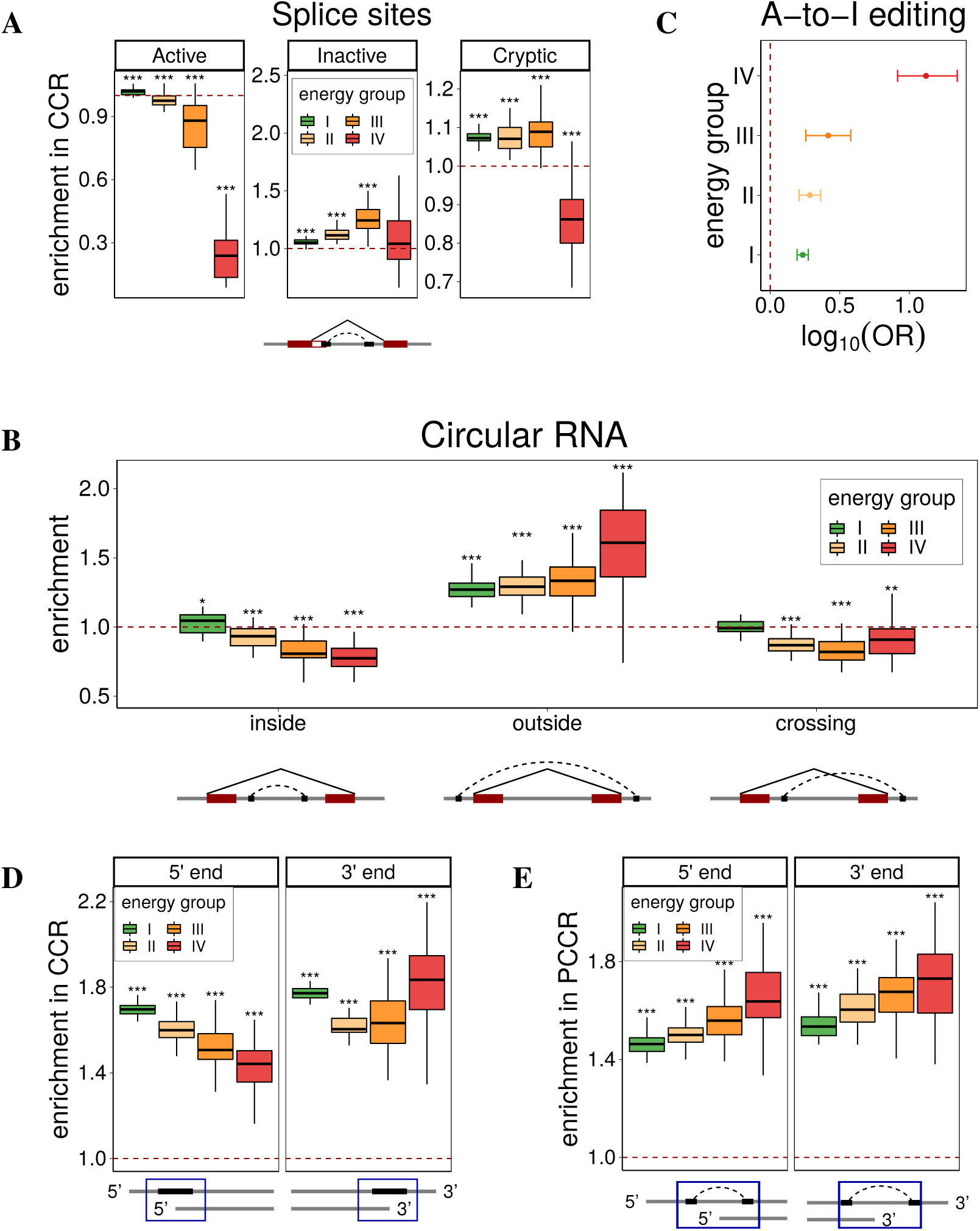
Splicing, RNA editing, and end processing. **(A)** CCRs are depleted around actively expressed splice sites and enriched around inactive and cryptic splice sites. **(B)** PCCRs are enriched outside of back-spliced introns (circular RNAs from TCSD, (*44*)) and depleted in inside and crossing configurations. **(C)** CCRs are enriched with A-to-I RNA editing sites (RADAR REDIportal (*50, 51*)); OR denotes the odds ratio (see Methods). **(D)** CCRs are enriched with 5’- and 3’-ends of transcripts annotated in GENCODE database (including all aberrant and incomplete transcripts). That is, transcript ends frequently occur in double-stranded parts of PCCRs. **(E)** PCCRs are also strongly enriched with 5’- and 3’-ends of transcripts, i.e. the annotated transcript ends frequently occur in the loop between double-stranded parts of PCCRs.

RNA secondary structure has been suggested to be a defining feature that leads to backsplicing and formation of circular RNAs (circRNA) (*43*). A comparison with the set of circRNAs from the tissue-specific circular RNA database (TCSD) (*44*) by random shift and random gene controls revealed a pattern opposite to that of linear introns, in which PCCRs were enriched outside circRNAs and depleted inside circRNAs (Figure 4B). This supports the hypothesis that circRNAs originate from loop-out sequences that are formed by stable double-stranded RNA structures (*43*).

### RNA editing

A widespread form of post-transcriptional RNA modification is the enzymatic conversion of adenosine nucleosides to inosine, a process called A-to-I editing, which is mediated by the ADAR family of enzymes (*45, 46*). Since ADAR editing occurs in double-stranded RNA substrates (*47, 48*), we asked whether A-to-I editing sites are enriched among CCRs (Figure 4C). We observed a strong enrichment of ADAR edited sites that are documented in the RADAR database (2,576,459 sites) within CCR as compared to non-CCR parts of conserved intronic regions (OR = 2.1 ± 0.2). While the odds ratio was nearly the same for PCCR with energy from −15 to −30 kcal/mol, it dramatically increased for PCCR with the energy below −30 kcal/mol (2.1±0.2 vs 13.2±5.0), reconfirming previous observations that ADAR targets are long double-stranded RNAs (*49*). The odds ratio was also significantly greater (1.7 ± 0.2 vs. 22.2 ± 13.1) for PCCR with higher evidence of compensatory mutations, but it didn’t depend significantly on the PCCR spread. These results were concordant with each other for the datasets of A-to-I editing sites from RADAR and REDIportal databases (*50, 51*) and can be regarded as an additional support for double-stranded structure of PCCRs.

### 5’-end and 3’-end processing

Recent reports indicate that RNA structure is important for the 3’-end processing of human mRNAs (*8*), and that competing RNA base pairings could be involved in alternative splicing and polyadenylation in the 3’-variable region (*52*). In this section we characterize the relationship between PCCRs, 5’-end, and 3’-end mRNA processing using the data from transcript annotation databases (*53*) and clusters of poly(A)-seq and CAGE-tags (*54, 55*).

First, we estimated the number of transcripts that start or end within CCRs, including all incomplete and aberrant isoforms that are annotated in GENCODE (Figure 4D). Both 5’- and 3’-ends were enriched inside CCRs suggesting that double-stranded structures are associated with suppression of aberrant transcripts. Next, we asked whether the annotated 5’- and 3’-ends of transcripts were enriched in the inner part of PCCRs, i.e. in the regions between paired CCRs. Indeed, both were significantly enriched, and the magnitude of the enrichment increased with increasing PCCR stability (Figure 4E). The random gene control confirmed that the effect was not due to non-uniform distribution of PCCRs. To rule out the possibility that non-expressed isoforms contribute to the observed enrichment, we additionally examined the expressed CAGE-tags and poly(A)-seq clusters and found that they were also significantly enriched within PCCR (Figure S9). Since PCCRs are, in turn, enriched inside introns, this motivated us to analyze triple associations between RNA structure, splicing, and end processing.

Thus, we asked whether the annotated transcript ends, CAGE-tags, and poly(A)-seq clusters occur more frequently in introns that contain PCCRs. However, introns with PCCRs tend to be also longer than introns without PCCRs, which may affect the above frequencies. Hence, we subdivided all annotated introns into two groups, introns with PCCR (IWP) and introns without PCCR (IWO) and selected two samples from IWP and IWO, 40,061 introns each, with matching intron lengths (Figure S10). 11,313 (33.1%) of IWP contained at least one annotated 3’-end, while only 11,031 (24.0%) of IWO did so (OR = 1.56 ± 0.05). Similarly, 10,631 IWP (31.1%) contained at least one annotated 5’-end, compared to 10,381 (22.6%) for IWO (OR = 1.54 ± 0.05). The enrichment of both 5’- and 3’-ends in IWP raises an intriguing hypothesis that there could be an RNA structure-mediated coupling between splicing and end processing (see Discussion).

### Mutations

Mutations generally lower the stability of RNA secondary structure, and some of them are under evolutionary selection owing to their effects on the thermodynamic stability of pre-mRNA (*56*). In order to estimate the impact of human population polymorphisms on PCCRs, we analyzed SNPs from the 1000 Genomes project (*57*) and compared SNP density in CCR with that in the remaining parts of CIR. Germline SNPs were significantly underrepresented in CCR (0.0196 vs 0.0207 SNPs per nt, P-value < 0.01). Next, for each SNP that occurs in a PCCR, we calculated the free energy change caused by the mutation and compared it to the free energy change that would have been observed if the same mutation occurred at a different position of the same CCR (Figure S11A). It turned out that actual SNPs destabilized PCCRs less than it would be expected by chance (Wilcoxon test, P-value < 0.01) suggesting that SNPs generally tend to minimize their destabilizing impact on RNA structure.

Compensatory mutations may also occur in the human population, but at a substantially lower frequency. To estimate the frequency of compensatory population polymorphisms, we identified PCCRs that contain SNPs with allele frequency of 1% and higher. Out of 64,074 base pairs in PCCRs that were affected by SNPs, only 12 showed sufficient support for compensatory mutations. After randomization, in which the pairs of complementary nucleotides were randomly switched, the respective average values were 64,132 and 0.06 (see Methods, Figure S11B). The density of compensatory mutations normalized to the number of base pairs with mutations were 0.019% for the actual base pairs and (9.0 ± 7.3) ∗ 10^−5^% for the randomized set. From this we conclude that, despite compensatory mutations within PCCR are quite rare in the human population, they are significantly enriched (Poisson test, P < 0.01).

### RBP binding sites

Multiple lines of evidence indicate that binding of RNA-binding proteins (RBPs), which is crucial for co- and post-transcriptional RNA processing, depends on RNA structural context (*58–60*). To assess the association between long-range RNA structure represented by PCCRs and RBP binding, we computed eCLIP peak frequencies (*61, 62*) in the vicinity of CCRs and compared them to the background eCLIP peak frequencies in conserved intronic regions surrounding CCRs. Among factors that showed a substantial enrichment there were RBPs that are known to exert their function in conjunction with RNA structure such as *RBFOX2* (*32*) and factors that favor increased structure over their motifs such as *SRSF9* and *SFPQ* (*63, 64*) (Figure 5A). For RBPs with single-stranded RNA binding activity such as *ILF3* and *hnRNPA1*, we observed a significant depletion of binding sites within CCRs (Figure S12) (*65, 66*).

**Figure 5:**
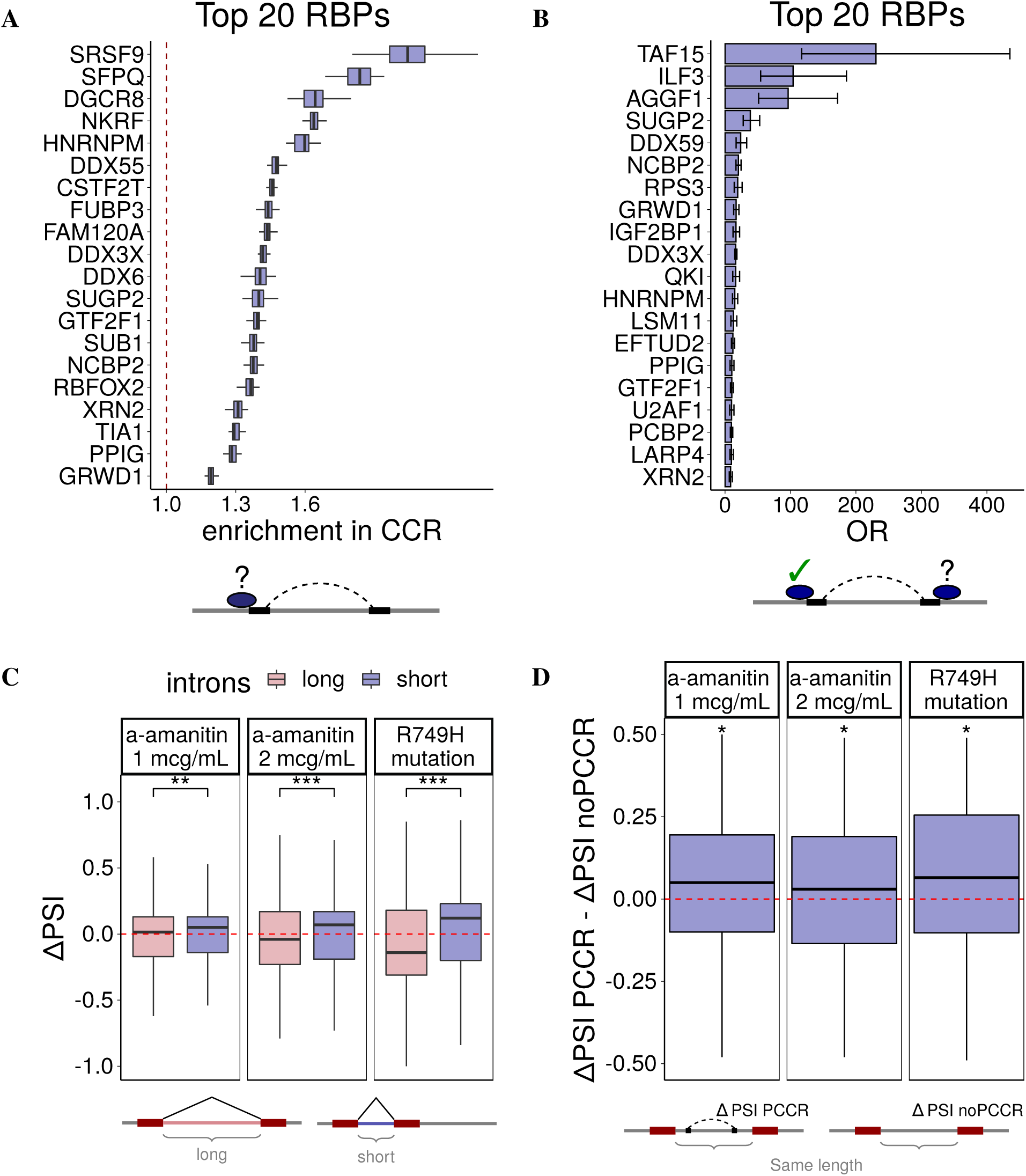
RNA-binding proteins (RBP). **(A)** According to eCLIP profiles, CCRs are enriched within binding sites of some RBPs (top 20 RBPs are shown). The RBPs that show depletion of CCRs are listed in Figure S12. Boxplots represent *n* = 40 random shifts of CCR within CIR. **(B)** The odds ratios (OR) of RBP binding near both CCR in PCCR given that RBP binds near at least one CCR indicate that PCCRs are enriched with forked eCLIP peaks. Boxplots represent *n* = 40 random shifts of CCR within CIR. **(C)** The change of inclusion rate (∆Ψ) of exons following short introns vs. exons following long introns in response to RNA Pol II slowdown with *α*-amanitin and in slow RNA Pol II mutants (*68*). **(D)** The difference between the inclusion rate change of exons following introns with a PCCR (∆ΨPCCR) and the inclusion rate change of exons following introns of the same length, but without PCCRs (∆ΨnoPCCR) in response to RNA Pol II slowdown.

One of the features of the eCLIP protocol is that an RBP can crosslink with either RNA strand that is adjacent to the double-stranded region. Hence, we expect a higher chance of observing an eCLIP peak near the CCR given that the other CCR in the pair contains a nearby peak, a situation that will be referred to as *forked* eCLIP peaks. To estimate the magnitude of this association, we computed the respective odds ratio and found that the vast majority of RBPs (64 out of 74, p-value < 10^−12^) indeed have a substantially higher likelihood of binding close to a CCR given that they bind the other CCR in a pair (Figure 5B). This can be regarded as independent evidence for double-stranded structure of PCCRs. Interestingly, the largest odds ratio was observed for *TAF15*, a TBP-associated factor 15, which is not a dsRNA binding protein itself, but interacts with FUS, which is capable of binding dsRNA (*67*). Similarly, *ILF3*, which showed a depleted RNA binding within CCRs, nevertheless is positively associated with forked eCLIP peaks, suggesting its binding at single-stranded regions adjacent to PCCRs. This indicates, on one hand, that RBP binding is inseparable from the surrounding RNA structure and, on the other hand, that eCLIP peaks may not correctly reflect the actual binding positions of RBPs since they are affected by intramolecular base pairings and interactions with other players.

### RNA Pol II elongation speed

The kinetic model of cotranscriptional splicing suggests that RNA Pol II elongation slowdown expands the “window of opportunity” for the recognition of weak splice sites, thereby increasing the rate of inclusion of upstream exons (*68, 69*). Besides this direct impact on splice site recognition, slow RNA Pol II elongation may also affect the way the transcript folds, which is another important determinant of how the transcript will be processed by the splicing machinery (*70*). To investigate the role of long-range RNA structure in co-transcriptional splicing, we performed RNA-seq experiments, in which we used *α*-amanitin to slow down the RNA Pol II elongation speed (*71*), and additionally analyzed publicly available data on the impact of RNA Pol II elongation speed on splicing (*68*).

The expected consequence of the RNA Pol II slowdown is that the inclusion rate of exons that follow short introns will increase, and the inclusion rate of exons that follow long introns will decrease. Indeed, this trend was observed both when RNA Pol II elongation speed was decreased by *α*-amanitin and in the slow RNA Pol II mutant R749H (Figure 5C) (*68*). To check whether RNA Pol II slowdown differently affects introns with and without PCCRs, we matched each exon that follows an intron containing a PCCR with a randomly chosen exon that follows an intron of the same length, but without PCCRs. The difference in inclusion rates of these matched exons showed that exons that follow an intron with a PCCR tend to be more included than exons following an intron without PCCRs at both concentrations of the inhibitor and in R749H RNA Pol II mutant (Figure 5D). This can be considered as evidence for RNA Pol II slowdown to affect exon inclusion through pre-mRNA folding, in addition to modulation of splice site recognition. Namely, slower RNA Pol II elongation speed may not only facilitate processing of upstream splice sites by the spliceosome, but also allow sufficient time for the intronic RNA structure to fold, thus promoting exon inclusion. A particular example of such kinetic mechanism linked to RNA structure was reported for the *Ate1* gene, in which a long-range base pairing dynamically regulates the ratio of mutually exclusive exons (*36*).

### Case studies

In this section, we focus on the association of long-range base pairings with RBP binding in the context of two particular splicing-related mechanisms, RNA bridges (*32*) and exon loop-outs (*33*).

To identify potential RNA bridges, i.e., long-range RNA structures that bring an RBP binding site closer to the regulated exon (*32*), we searched for candidate binding sites of RBPs profiled by eCLIP that were located within 50 nts from a CCR, on one hand, and exons located within 50 nts from its mate CCR, on the other hand. To detect regulation, we additionally required that exon inclusion rate significantly respond to shRNA-KD of the same factor using the data on RBP knockdowns produced by the ENCODE Consortium (see Methods) (*62*). This procedure yielded a set of 296 candidate RNA bridges (Supplementary Data File 3), including the RNA bridge that controls the inclusion of exon 12 of *ENAH* gene (Figure 6A). A PCCR with the hybridization energy −19.8 kcal/mol coincides the core part of the RNA stem that was reported earlier (*32*). We reconfirm that it is surrounded by forked eCLIP peaks of *RBFOX2*, which reflects cross-linking next to the double-stranded region, and that the inclusion of exon 12 drops by 43% (∆Ψ = −0.43) upon *RBFOX2* knockdown. However, we also find a nested PCCR with the hybridization energy −20.4 kcal/mol, which suggests that the RNA bridge extends much further than it was reported originally. As a novel example, we describe a candidate RNA bridge in the 3’-end of *RALGAPA1* gene, which encodes the major subunit of the RAL-GTPase activating protein (*72*). In this gene, a group of nested PCCRs approximates binding sites of *RBFOX2* and *QKI* to the penultimate exon (Figure 6B). The knockdown of each of these factors promotes exon skipping, which indicates that exon inclusion depends on the binding of *RBFOX2* and *QKI* through an RNA bridge.

**Figure 6:**
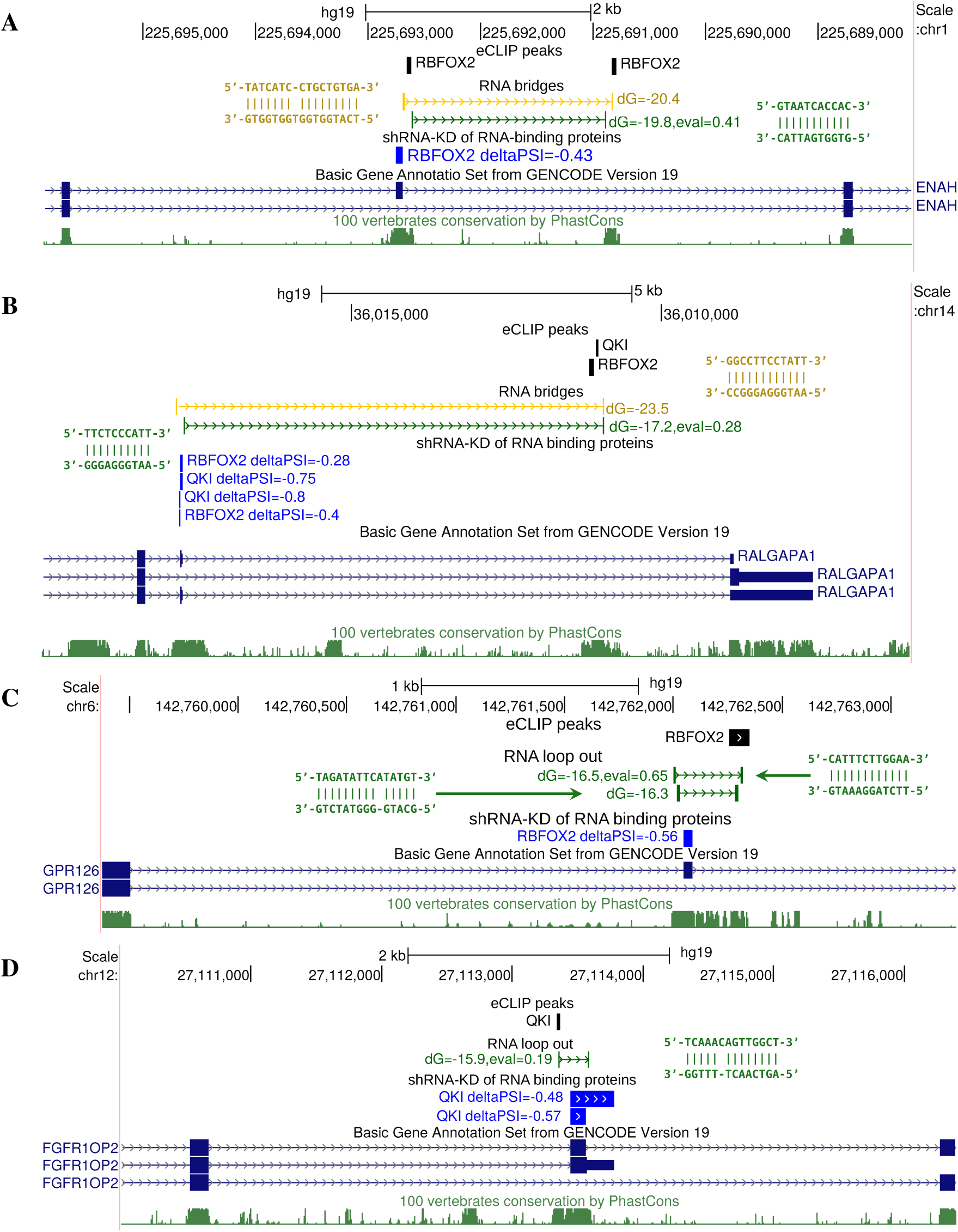
Case studies. **(A)** An RNA bridge in *ENAH* gene brings a distant RBFOX2 binding site into proximity of the regulated cassette exon (*32*). The exon inclusion rate substantially decreases under RBFOX2 depletion (∆Ψ = −0.43). **(B)** The predicted RNA bridge in *RALGAPA1* brings distant binding sites of RBFOX2 and QKI to the regulated exon. The exon significantly responds to the depletion of these two factors (∆Ψ = −0.28 and ∆Ψ = −0.75, respectively). **(C)** A cassette exon in *GPR126* is looped out by a PCCR overlapping an eCLIP peak of RBFOX2 and significantly responds to RBFOX2 depletion (∆Ψ = −0.56). **(D)** An alternative terminal exon in FGFR1OP2 is looped out by a PCCR overlapping an eCLIP peak of *QKI* and significantly responds to *QKI* depletion (∆Ψ = −0.48). In all panels, exon inclusion rate changes are statistically significant (q-value < 0.01).

In a similar way, we identified candidate base pairings that loop out exons by searching for PCCRs that surround an exon and contain an RBP binding site within one of the CCRs. Additionally, we required that the exon significantly respond to RBP knockdown. This procedure yielded a set of 1135 candidate exon loop-outs (Supplementary Data File 4). Among them there were two nested RNA structures looping-out exon 24 of *GPR126*, the human G protein-coupled receptor 126, in which one of the CCRs overlaps a *RBFOX2* binding site, and the exon responds to *RBFOX2* knockdown (Figure 6C). Another example is the alternative 3’-end exon in *FGFR1OP2*, Fibroblast Growth Factor Receptor 1 Oncogene Partner 2, which is suppressed by *QKI* knockdown and, at the same time, is looped out by a PCCR that overlaps a *QKI* binding site (Figure 6D).

### Data visualization and availability

The tables listing PCCRs for GRCh37 and GRCh38 human genome assemblies are available in BED format as Supplementary Data Files 1 and 2, respectively. These predictions are visualized through a track hub for the UCSC Genome Browser (*31*). In order to connect the track hub, the following link https://raw.githubusercontent.com/kalmSveta/PCCR/master/hub.txt can be copied and pasted into the form https://genome.ucsc.edu/cgi-bin/hgHubConnect#unlistedHubs. The PCCR tracks are grouped by the energy groups, spread, and E-value. Along with PCCRs, we additionally report the response of exons to RNA Pol II elongation slowdown, icSHAPE reactivity scores, and RIC-seq predictions. The tables listing the predicted RNA bridges and looping-out PCCRs are available as Supplementary Data Files 3 and 4, respectively. They are visualized through (https://raw.githubusercontent.com/kalmSveta/RNA-bridges/master/hub.txt) along with eCLIP data and exon responses to shRNA knockdowns. All Supplementary Data Files are available online at http://arkuda.skoltech.ru/~dp/shared/PrePH/.

## Discussion

It has been increasingly acknowledged that RNA structure plays a critical role in the regulation of eukaryotic gene expression at all steps from transcription to translation, but very little attention has been paid to long-range RNA structure. From the thermodynamic standpoint, long-range base pairings contribute to the enthalpy of RNA folding as much as local base pairings do, and the corresponding energy figures exceed by an order of magnitude the typical folding energies of globular protein domains (*73*). However, since RNA structure affects, and is itself strongly affected by RBP binding, a reliable prediction of long-range RNA structure in full-length eukaryotic transcripts doesn’t seem feasible at the current state of the art. Instead of the detailed structure, here we consider as a proxy for RNA structure the core of highly stable and evolutionarily conserved double-stranded regions, different combinations of which may represent one or several physiologically relevant folds.

The existence of associations between long-range RNA structure and splicing has been noted in previous studies (*7*), including our earlier reports (*27–29*). The trends that were proposed for smaller sets also hold for the extended catalog of PCCRs presented here, namely the preference of PCCRs to be positioned within introns proximally to splice sites (*27*), lower inclusion rate of looped-out exons (*33, 74*), circumscription of circRNAs (*75*), avoidance of intronic branch points (*76, 77*), and generally obstructive effect on splice sites that are implicated in double-stranded structure (*78–80*). We additionally observed a remarkable overlap of PCCRs with A-to-I RNA editing sites and multiple associations with forked eCLIP peaks, which reveal traces of multi-molecular complexes with patterns that are specific to double-stranded regions. These new findings reconfirm the well-known mechanism of ADAR-mediated pathway (*81*) and show the importance of RNA structure for the assembly of RNA-protein complexes, with preference for some RBPs and avoidance for the others (*62, 63*). At the same time, they may also be regarded as independent support for double-stranded structure of PCCRs in addition to the evidence from experimental RNA structure profiling and compensatory substitutions.

The components of RNA processing machinery operate in a strict coordination not only in space but also in time. The kinetic profile of RNA Pol II elongation has a significant impact on alternative splicing (*82*). Slow RNA Pol II elongation generally opens a window of opportunity for weak splice sites to be recognized, leading to higher inclusion of alternative exons, although in some cases the effect can be quite opposite (*83, 84*). Slow RNA Pol II elongation may also influence poly(A) site choice by enhancing the recognition of suboptimal polyadenylation signals (*85*). Consistent with this, we observe an increased inclusion of exons that are preceded by shorter introns. However, we also observe an additional component to this general trend, one in which structured and unstructured RNAs respond differently to RNA Pol II slowdown. This observation indicates that long-range RNA structure could coordinate the interaction between spatial and temporal components of splicing regulation. The example of long-range RNA structure in the *Ate1* gene demonstrates that mechanisms similar to bacterial attenuation may also take place in eukaryotic cells (*36, 86, 87*).

RNA structure is implicated in the recognition of polyadenylation signals (PAS) by cleavage and polyadenylation specificity factor (CPSF) and facilitates the 3’-end processing by juxtaposing PAS and cleavage sites that are otherwise too far apart (*8, 88*). Functional RNA structures in 5’-UTRs are also implicated in the regulation of translation (*11*). While these reports mostly concern local RNA structure, an intriguing finding of this work is the link between long-range RNA structure and pre-mRNA 3’-end processing, which is manifested by the enrichment of poly(A)-seq clusters in the inner part of PCCRs. It indicates that mechanisms other than sequestration or spatial convergence of PAS and cleavage sites may be involved (*89, 90*). In fact, human genes contain thousands of dormant intronic PASs that are suppressed, at least in part, by U1 small nuclear ribonucleoproteins in a process called telescripting (*91*). While the exact mechanism of this suppression is not known, many intronic PASs were found to be associated with *CstF64*, a ubiquitous pre-mRNA 3’-processing factor (*92*), suggesting that cleavage and polyadenylation machinery may actually operate in all introns constitutively. Could it be that RNA structure helps to suppress premature intronic polyadenylation?

Figure 7 illustrates a hypothetical mechanism of suppression of intronic polyadenylation by co-transcriptional splicing, which explains the enrichment of poly(A)-seq clusters in the inner parts of PCCRs. Indeed, the cleavage and polyadenylation of a structured pre-mRNA could be rescued by co-transcriptional excision of the intron while RNA structure stabilizes the molecule through intramolecular base pairings despite disruption of the backbone (Figure 7A). However, such a rescue won’t happen in unstructured RNAs when splicing has a delay relative to cleavage and polyadenylation (Figure 7B). This scenario is further supported by a conspicuous association between transcript 5’- and 3’-ends in exhaustive transcriptome annotations, a typical example of which is an intron that contains a 3’-end of protein-coding transcript and a 5’-end of another, usually non-coding transcript (Figure S7). It is 2.86 ± 0.10 times more likely to see a 5’-end in an intron that contains a 3’-end, and hence the enrichment of poly(A)-seq clusters within PCCRs also implies the enrichment of 5’-ends and CAGE clusters. The described mechanism could be responsible for the generation of transcripts with alternative 3’-ends, e.g., for the RNA structure-mediated switch between splicing and polyadenylation in the *Nmnat* gene in *D. melanogaster* (*27*).

**Figure 7:**
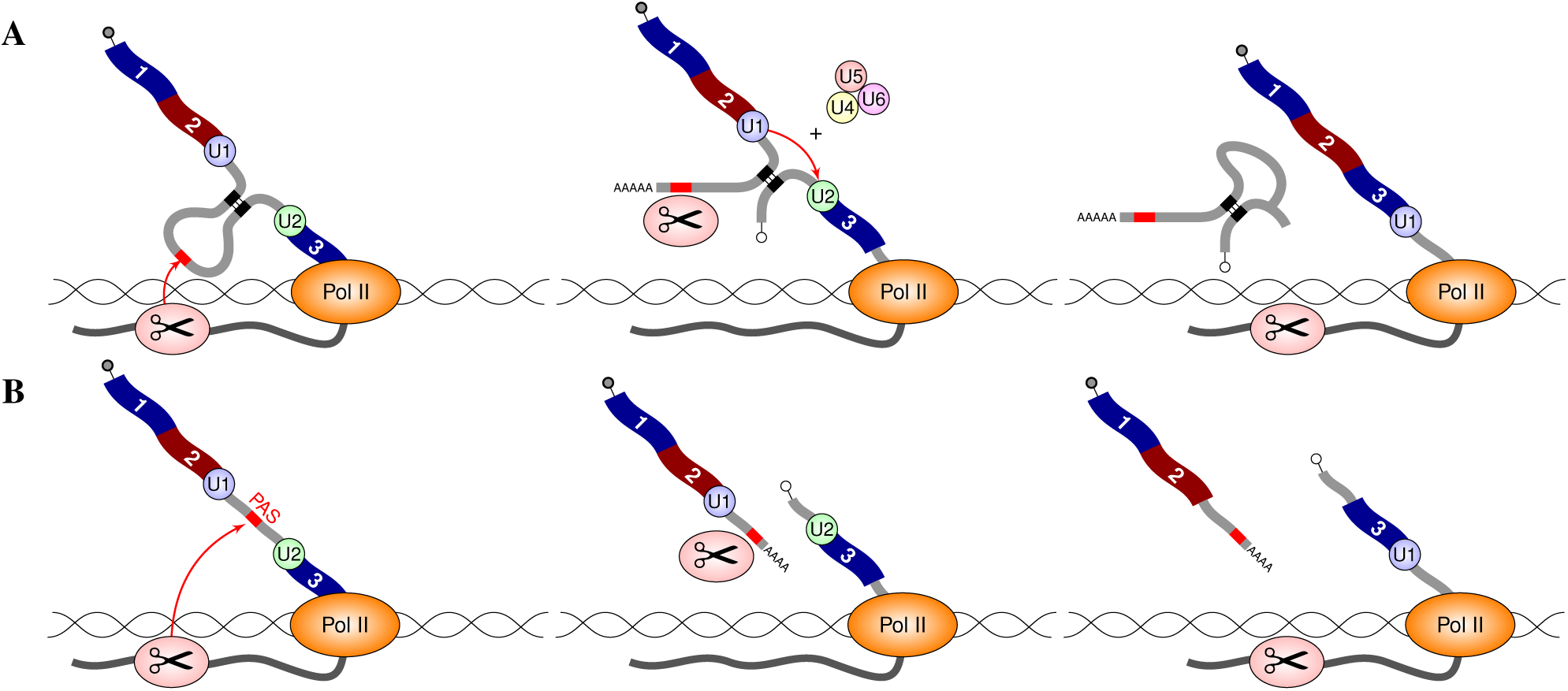
RNA folding and splicing could mediate co-transcriptional suppression of premature cleavage and polyadenylation (a hypothesis). **(A)** The cleavage and polyadenylation of a structured pre-mRNA is rescued by the co-transcriptional excision of the intron while RNA structure stabilizes the molecule through intramolecular base pairings. **(B)** In the absence of RNA structure, such a rescue won’t happen when splicing has a delay relative to cleavage and polyadenylation. Switching between (A) and (B) depends on the rates of splicing, folding, and Pol II elongation.

RNA structure probing by icSHAPE and the assessment of long-range RNA–RNA interactions by photo-inducible RNA crosslinking are the most current techniques for global analysis of RNA structure *in vivo* (*18, 22–24*). However, these data reflect gene expression patterns that are specific to cell lines, in which they were generated, and have strong undercoverage bias in intronic regions, which are spliced out and degraded. The reduction of the intronic signal is also a common problem for RNA crosslinking and immunoprecipitation experiments (*93*). Additionally, it was meaningful to compare PCCRs only to intramolecular RNA contacts that belong to conserved intronic regions and don’t exceed the distance limit of 10,000 nts. Consequently, the validation with respect to these assays was possible only for a small number of PCCRs, on which the comparison, however, showed a concordant result (Table 2). On the other hand, the fact that 2,961 out of 916,360 PCCRs were supported by RIC-seq is highly significant compared to the expected intersection of 381 structures for two interval sets of the same size obtained by random shifts.

The major problem of the current method is the high number of false positive predictions. On the one hand, the procedure for the estimation of FDR is based on the assumption that pre-mRNAs of different genes do not interact with each other, and that regulatory sequences in different genes evolve independently. However, psoralen crosslinking and proximity RNA ligation assays have demonstrated that this assumption may not be completely true because RNA-RNA interactions *in trans* are very abundant (*20–23*). Therefore, the rewiring control overestimates FDR, as it did in previous works (*29*). On the other hand, the amount of random complementarity in conserved intronic regions is, indeed, very large. For instance, two intronic binding sites of a transcription factor that occur on opposite DNA strands will be detected as a PCCR, and they may even be supported by spurious compensatory substitutions that result not from selective constraints on RNA structure, but from evolving specificity of the binding site. An example of this is the RP11-439A17.4 lncRNA, a part of which is complementary to conserved sequences in 22 mammalian histone genes, but the complementary elements are, in fact, the binding sites of MEF-2A, a myocyte-specific enhancer factor (*29*). The evolution maintains them conserved and technically complementary for reasons other than base pairing, which makes this situation in principle indistinguishable from evolutionary selection acting on true RNA structures. To reduce FDR, a significant improvement could be achieved by combining the methodology presented here with the emerging experimental strategies for profiling of RNA–RNA contacts such as RIC-seq (*24*). This appears to be the most promising direction for future research.

## Conclusion

Eukaryotic pre-mRNAs are structured macromolecules with base pairings that span long distances, and RNA secondary structure plays a critical role in their processing. Here, we present the most complete to-date catalog of conserved complementary regions in human protein-coding genes and provide their extensive characterization. In spite of high false positive rate, the predicted double-stranded RNA structures show significant associations with virtually all steps of pre-mRNA processing. We offer this catalog for common use as a reference set and provide its convenient visualization through a UCSC Genome Browser track hub.

## Acknowledgements

RNA-sequencing was funded by Skolkovo Institute of Science and Technology. The authors would like to thank Skoltech Genomics Core facility staff (Margarita Ezhova, Maria Logacheva) for library preparation and sequencing, and the `Cobrain` computer cluster for access to computational facilities. We also thank Prof. Yuanchao Xue and Prof. Changchang Cao for providing the list of clusters of chimeric reads for intramolecular interactions. S.K., S.D. and D.P. were supported by Russian Foundation for Basic Research grant 18-29-13020-MK and by Skolkovo Institute of Science Technology Research Grant RF-0000000653. M.K. and D.S. were supported by Russian Foundation for Basic Research grant 18-29-13020-MK.

## Author contributions

D.P. and S.K. designed and carried out the analysis. M.K. and D.S. performed the experiments. S.D. and A.M. analyzed the evolutionary part. All authors analyzed the data and discussed the results. D.P. and S.K. wrote the paper.

## Methods

### Genomes and transcript annotations

February 2009 (hg19, GRCh37) and December 2013 (hg38, GRCh38) assemblies of the human genome were downloaded from Genome Reference Consortium (*94*). These assemblies were used with GENCODE transcript annotations v19 and v33, respectively (*95*). Only genes labelled “protein coding” were analyzed. Transcript annotations were parsed by custom scripts to extract the coordinates of exons and introns. The results for GRCh37 assembly are reported throughout the paper, however Genome Browser track hubs (see below) contain tracks for both GRCh37 and GRCh38 assemblies. The intronic regions were defined as the longest continuous segments within genes which don’t overlap any annotated exons (including exons of other genes). The intronic regions were extended by 10 nts into the flanking exons to enable identification of CCRs that overlap splice sites such as regions R1–R5 in the human *Ate1* gene (*36*). Next, these regions were intersected with the set of conserved RNA elements (phastCons elements track for the alignment of 99 vertebrates genomes to the human genome (*96*)) using bedtools (*97*).

### PrePH

The first reference to long-range RNA structure appeared in the literature under the term “pan-handle structure”, which was coined in by virologists in the 1980’s to refer to a complementary base pairing between the 5’-end and 3’-end of the RNA genome of several segmented negative-stranded viruses (*98*).

The PrePH (PREdiction of PanHandles) utility uses a *k*-mer-based technique, which is similar to previously published IRBIS method (*29*), to identify all pairs of nearly-perfect complementary regions in a given pair of sequences. At the preparatory step, PrePH pre-computes a 4^*k*^ × 4^*k*^ table containing helix hybridization energies for all pairs of *k*-mers (default *k* = 5) that are either Watson-Crick complementary, or contain a few GT base-pairs (by default at most two) using energy tables from Vienna RNA package (*99*). The dynamic programming matrix is computed by local Smith-Waterman algorithm using a limited set of structural rules: initiating a *k*-nt-long helix, extending a helix by a stacking base pair, and adding to a helix a short internal loop or bulge with up to *m* nucleotides in each strand (default *m* = 2), followed by another helix. To speed up backtracking, PrePH uses auxiliary matrices to store intermediate structures, and reports non-intersecting pairs of complementary regions passing the energy threshold (by default −15kcal/mol). Here, two pairs of complementary regions, *x*_1_ complementary to *y*_1_ and *x*_2_ complementary to *y*_2_, are referred to as intersecting if *x*_1_ has common nucleotides with *x*_2_ and also *y*_1_ has common nucleotides with *y*_2_. That is, two pairs of complementary regions may intersect by only one, not both interacting strands. The reduced scoring scheme, optimized back-tracking, and indexing of the initial sequences by *k*-mers result in a great improvement of computation speed. The detailed description of PrePH is exempt to Supplementary Methods. PrePH software is available at github (https://github.com/kalmSveta/PrePH).

### Benchmark

We compared the accuracy and runtime of PrePH to those of other programs such as IntaRNA2.0 (*100*), RIsearch2 (*101*), RNAplex (*102*), DuplexFold (*103*) and bifold (*103*). PrePH was run with the following parameters: *k*-mer length is 5 nts, maximal distance between complementary regions is 10,000 nts, the minimal length of the aligned regions is 10 nts, the energy threshold is −15 kcal/mol, the maximal number of Wobble pairs in a *k*-mer is 2. The parameters for the other programs are listed below.

To benchmark the time efficiency, we use a set of 1000 pairs of randomly chosen conserved intronic sequences from the human genome. The sequences were 50 to 500 nucleotides-long and contained nearly perfect sequence complementarity. All the programs were run with the energy threshold set to −15 kcal/mol. `IntaRNA2.0` *outOverlap* parameter was set to *B*, which allowed overlap for interacting subsequences for both target and query; *n* parameter was set to 100 to limit the maximal number of suboptimal structures; *qAcc* and *tAcc* were set to N to omit the computation of accessibility. `RIsearch2` seed length was set to 5, the length of flanking sequences considered for seed extension was set to 50. `RNAplex` *fast-folding* parameter was set to *f2* to allow the structure to be computed based on the approximated model. `DuplexFold` and `bifold` maximum loop/bulge size was set to two. All other parameters were left at their default values. The computations were carried out on Intel R Core TM i5-8250U CPU with 1.60 GHz. PrePH showed the quickest result compared to the other programs (191.4 sec) (Table S2A). At that, the equilibrium free energies of the predictions by PrePH correlated reasonably well with those of RNAplex, IntaRNA, and Duplexfold (Figure S13).

For the comparison of MFE between different methods, we used simulated data with 1000 pairs of nearly perfect sequence complementarity, which were from 10 to 50 nts. All the programs were run with the energy threshold set to −15 kcal/mol. The other parameters were as before. Pearson correlation coefficients were computed between energies of the predicted optimal structures. To compare the predictions of PrePH with predictions of other programs at the level of individual base pairs, we computed the following metric

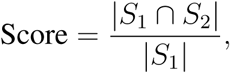

where *S*_1_ is the set of base pairs predicted by PrePH, *S*_2_ is the set of base pairs predicted by the other program, and *S*_1_ ∩ *S*_2_ is the common set of base pairs (|*S*| denotes cardinality of a set). The number of base pairs that were common between PrePH and each of the other programs as a fraction of the number of base pairs predicted by PrePH alone was used as a measure of specificity (Table S2B). PrePH showed the specificity above 80% with respect to all other programs except bifold, however the latter was not in agreement with all other programs.

We conclude that PrePH allows for computationally-efficient detection of PCCRs without significant loss of accuracy compared to other methods. The computation time of PrePH on the complete dataset of conserved intronic regions was 4 hrs (15 threads, 1200 MHz CPUs each).

### Relative position within the gene

The relative position of a genomic interval [*x, y*] in the containing gene [*a, b*], where *x, y, a*, and *b* are genomic coordinates on the plus strand, was calculated as 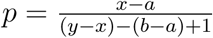 for the genes on the positive strand. For the genes on the negative strand the value of 1 − *p* was used instead.

### Gene Ontology (GO) enrichment analysis

Gene Ontology (GO) enrichment analysis was performed by controlling for the gene length since genes with PCCRs tend to be longer than genes without PCCRs. We randomly matched each gene with PCCRs to a gene without PCCR of approximately the same length. The enrichment of GO terms (*104, 105*) between these two gene sets was calculated with clusterProfiler R package (*106*).

### Experimental RNA structure data

The icSHAPE reactivity scores (*107*) were mapped from GRCh38 to GRCh37 human genome assembly by LiftOver (*108*). The reactivity score of a CCR was calculated as the average reactivity of its base paired nucleotides, for which the icSHAPE reactivity score was available. The background reactivity was calculated as the average reactivity of the same number of nucleotides chosen at random outside the CCR, but within its conserved intronic region. Wilcoxon signed rank test was applied to matched samples of reactivity score differences to test for departures from zero.

The coordinates of base paired regions from PARIS experiments (*22*) (15,036 pairs) were mapped from GRCh38 to GRCh37 by Liftover (*108*). The coordinates of base paired regions from LIGR-seq data (551,926 pairs) were used in the GRCh37 human genome assembly (*23*). The coordinates of intramolecular base-paired regions from RIC-seq data (*24*) (501,144 pairs) were kindly provided by Prof. Xue by request (Supplementary Data File 5). In all three datasets, we selected the interacting pairs that were located intramolecularly within CIR of protein-coding genes from 1 to 10,000 nt apart from each other. This resulted in 907 such pairs for PARIS, 586 for LIGR-seq, and 1,804 for RIC-seq. In order to evaluate the precision and recall, we selected CIR with at least one nucleotide overlapping the experimentally validated structures and confined our analysis to PCCRs located in these regions. A CCR was classified as a true positive if it had at least one common nucleotide with an experimentally validated structure. A PCCR was classified as a true positive if both its CCRs intersected by at least one nucleotide with an experimentally validated such pair.

### Cell culture, treatments and RNA purification

A549 cell line was maintained in DMEM/F-12 medium containing 10% fetal bovine serum, 50 U/ml penicillin, and 0.05 mg/ml streptomycin (all products from Thermo Fisher Scientific) at 37°C in 5% CO_2_. For *α*-amanitin (Sigma) treatments 1 and 2 *µ*g/mL of *α*-amanitin was added to cells at 50-70% confluency. After 24h of treatment, cells were harvested, total RNA was isolated using PureLink RNA Mini Kit (Thermo Fisher Scientific). Poly(A)^+^ mRNA was purified using Dynabeads Oligo(dT) 25 (Thermo Fisher Scientific) following the manufacturer’s instructions.

### Library preparation and RNA sequencing

Illumina cDNA libraries were constructed using NEBNext Ultra II Directional RNA Library Prep Kit for Illumina (New England BioLabs) following the manufacturer’s protocol with the only modification: the change of fragmentation time from 15 to 10 minutes. cDNA libraries were sequenced using the NextSeq500 (Illumina, San Diego, CA USA) instrument; 33–41 million raw reads were obtained for each sample with a 75 bp read length. The results of RNA-sequencing have been deposited at Gene Expression Omnibus under the accession number GSE153303.

### Splicing quantification

RNA-seq data of poly(A)^+^ RNA for the HepG2 cell line (accession numbers ENCFF670LIE and ENCFF074BOV) were downloaded in BAM format from the ENCODE Consortium website (*109*). Short-hairpin shRNA knock-down (shRNA-KD) of 250 RBPs followed by RNA-seq data (*62*) were downloaded in BAM format from ENCODE data repository (*109, 110*) (Table S3). Poly(A)^+^ RNA from wild-type *Amr* and *Rpb1* C4/R749H mutant HEK293 cells treated with *α*-amanitin for 42 h were downloaded from the Gene Expression Omnibus (GSE63375) (*68*). RNA-seq data of poly(A)^+^ RNA data from the A549 cell line treated with *α*-amanitin were obtained as explained below and mapped to the GRCh37 human genome assembly using STAR aligner with the default settings (*111*). The coordinates of circRNAs expressed in liver tissue and their associated SRPTM metrics (the number of circular reads per number of mapped reads per read length) were obtained from TCSD database (*44*). The genomic coordinates of adenine branch point nucleotides were selected from the validated set of branch points expressed in K562 cells (*39*).

RNA-seq experiments were processed by IPSA pipeline to obtain split read counts supporting splice junctions (*112*). Split read counts were filtered by the entropy content of the offset distribution, annotation status and canonical GT/AG dinucleotides at splice sites, and pooled between bioreplicates. The exon inclusion rate (Ψ, PSI, or Percent-Spliced-In) was calculated according to the equation

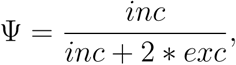

where *inc* is the number of reads supporting exon inclusion and *exc* is the number of reads supporting exon exclusion. Ψ values with the denominator below 10 were considered unreliable and discarded. Differential exon inclusion between a pair of conditions (shRNA-KD vs. non-specific control and *α*-amanitin vs. untreated control) was assessed as described previously (*113*).

Cryptic and actively-expressed splices sites in the human transcriptome were identified using genomic alignments of RNA-seq samples from the GTEx Consortium (*40*). Splice sites with the canonical GT/AG dinucleotides were called from split read alignments and ranked by the total number of supporting split reads pooled across all 8,551 samples. The top 2% (respectively, bottom 2%) of splice sites among those supported by at least three split reads were referred to as active (respectively, inactive). In order to identify cryptic splice sites, we applied the same strategy as (*114*) by scanning the intron sequences for any sites that have a MaxEntScan score >800 for donor sites and >950 for acceptor sites (*115*) and excluding splice sites that were detected in GTEx or present in the genome annotation. MaxEntScan score thresholds were chosen to have a comparable number of splice sites as in active and inactive sets above.

### 5’-end and 3’-end RNA processing

The genomic coordinates of human poly(A) sites profiled by high-throughput sequencing of 3’-ends of polyadenylated transcripts (poly(A)-seq) that were supported by 20 or more reads and intersected with the annotated transcript ends in GENCODE database were used (*54*). Similarly, clusters of human CAGE tags expressed in the HepG2 cell line that were supported by RPKM of at least 10 and intersected with the annotated transcript starts in GENCODE database were used (*116*).

### RNA editing

Adenosine-to-inosine (A-to-I) RNA editing sites were obtained from RADAR (*50*) and REDI-portal (*51*) databases. To compare the density of RNA editing sites in CCR and that in the adjacent conserved regions, we computed the number of adenosine residues that are RNA editing sites within CCR and compared it to the respective figures for conserved intronic regions of the same length outside of CCR. Odds ratio was calculated for the contingency table of adenosine residues that are/are not RNA editing sites within/outside CCR.

### eCLIP

Enhanced cross-linking and immunoprecipitation (eCLIP) peaks for 74 RBPs assayed in HepG2 human cell line were downloaded from the ENCODE data repository in bed format (*62, 109, 110*) (Table S4). The peaks were filtered by the conditions logFC ≥ 3 and p-value < 0.001. Since the agreement between two replicates was moderate, we use the pooled set of eCLIP peaks. To quantify the association between RBP binding and individual CCR, we calculated for each RBP the number of CCRs that intersected with its eCLIP peaks by at least 50% of the CCR length. This number was compared to the respective number of intersections obtained in the random shift control. To quantify the association of RBP binding with both CCRs in a PCCR, we constructed a contingency table for the number of PCCRs that had/didn’t have eCLIP peaks in left/right CCR for each RBP, and computed the respective odds ratio (OR). The 95% confidence interval for the odds ratio was calculated by Fisher test.

### Population polymorphisms

To evaluate the enrichment of population polymorphisms in CCR, we used genotyping data from phase 3 of the 1000 Genomes Project for 2504 individuals (*57*) and computed the density of SNPs (SNVs present in more than 0.1% of individuals) in stacked nucleotide pairs of CCR and compared it to the density of SNPs in conserved regions outside CCRs (regions of each type were merged with `bedtools merge` (*117*) before calculating the densities).

To assess the impact of SNPs on RNA structure stability, we calculated PCCR energy for the mutated sequence using energy parameters (*99*) and compared it to the respective energy of the structure with SNPs of the same substitution type as observed originally, but introduced at a different position. We selected PCCRs with the free energy change greater than 2 kcal/mol by absolute value and compared the two energy sets using Wilcoxon signed-rank test.

To evaluate the enrichment of compensatory mutations in PCCRs, we selected SNVs that occur in more than 1% of donors in the 1000 Genome Project and intersected their list with the list of base pairs in PCCRs. Among them we estimated the number of base pairs, in which a compensatory mutation occurred in more than 1% of donors. To estimate the expected number of base pairs with compensatory mutations, we first subdivided base pairs into groups composed of the same base pair types (AT, TA, CG, GC, TG, GT) located on the same chromosome with the same number of SNP donors. Then we randomly interchanged (“re-wired”) base pairs within each group, e.g. A_1_T_1_ and A_2_T_2_ were replaced by A_1_T_2_ and A_2_T_1_ and applied the same procedure again, i.e., estimated the number of base pairs, in which compensatory mutations defined by SNVs occurred in more than 1% of donors. (Figure S6A). This randomization procedure was repeated 100 times.

### Sequence conservation and complementary substitutions

To assess the degree of evolutionary conservation of a CCR, we computed the difference between the average PhastCons conservation score (*96*) of all its nucleotides and the average PhastCons conservation score of the same number of nucleotides in its flanking regions within the same `phastConsElements` interval.

To assess the number of complementary substitutions in PCCRs and their statistical significance, we used global multiple sequence alignments (MSA) of 99 vertebrate genomes with human genome (*118*). For each PCCR, we extracted two parts of the MSA corresponding to two CCRs using Bio.AlignIO.MafIO module from biopython library (*119*). The organisms that had indels compared to the reference organism (hg19) in any of the two CCRs were removed. The number of orthologous sequences for each PCCR ranged from 15 to 99. The two alignment blocks were merged through an additional spacer containing 10 adenine nucleotides, resulting in a MSA STOCKHOLM format with a secondary RNA structure generated by PrePH. Next, we restrict the phylogenetic tree for the original MSA (*118*) to have only the organisms available for the given PCCR and pass the tree and MSA to Rscape v1.2.3 (*37*) with the following parameters: *-E 1 -s –samplewc –nofigures*. The output .out files of Rscape were parsed by custom scripts to extract E-values of individual base pairs. The E-value of the PCCR was defined to be equal to the product of E-values of the base pairs that were marked as having significant covariations by R-scape. As a result of this procedure, E-values were obtained for 909,146 PCCRs; 539,264 E-values for PCCRs that were less than 1 were adjusted using Benjamini-Hochberg correction. MSA and the phylogenetic trees were downloaded from the UCSC Genome Browser website (http://hgdownload.cse.ucsc.edu/goldenpath/hg19/multiz100way/). Structural alignments were visualized using tableGrob function from gridExtra R package.

### Statistical analysis

The data were analyzed and visualized using R statistics software version 3.4.1 and ggplot2 package. Non-parametric tests were performed by built-in R functions using normal approximation with continuity correction. MW denotes Mann-Whitney sum of ranks test. Error bars in all figures and the numbers after the ± sign represent 95% confidence intervals. One-sided P-values are reported throughout the paper. The levels of significance 0.05,0.01,0.001 in all figures are denoted by ^∗, ∗∗^, and ^∗∗∗^, respectively.

## Data availability

The RNA-seq data generated in this study are available through Gene Expression Omnibus under the accession number GSE153303. Other data that were analyzed in this study are available via references listed in Methods section. The results of this study are available online at http://arkuda.skoltech.ru/~dp/shared/PrePH/.

## Code availability

The software developed in this study is available at https://github.com/kalmSveta/PrePH.

