## Supplementary information for "Conserved long-range base pairings are associated with pre-mRNA processing of human genes"

### Supplementary Methods

#### PrePH

The algorithm of PrePH (the PREdiction of PanHandles), which borrowed some essential ideas from its predecessor IRBIS (*29*), and can be considered as a sparse implementation of Smith-Waterman algorithm for local sequence alignment (*121*) with specific enhancements that look for complementarity of $k$-mers. The main feature that enables efficient computations on the genome-wide scale is that PrePH scoring scheme is based on $k$-mers instead of single nucleotides, i.e., we disregard structural units that contain less than $k$ base-paired nucleotides in a row. In order to compute the free energy contribution, we consider only a limited set of structural elements (stacked base pairs, bulges, and interior loops), neglect contributions from dangling ends and terminal mismatches, and disregard branching structures. Given a pair of sequences of lengths $M$ and $N$, we sequentially compute three dynamic programming matrices $D_{ij}$, $B_{ij}$, and $S_{ij}$ of size $M \times N$ as follows. We denote by $D_{ij}$ the minimum energy (the maximum score) of the interaction that ends at the position $i$ in the first sequence and the position $j$ in the second sequence. We denote by $B_{ij}$ the matrix that stores start coordinates of the base pairing that ends at the positions $i$ and $j$, while the matrix $S_{ij}$ stores the structure of the interaction in dot-bracket notation. The matrices $B_{ij}$ and $S_{ij}$ are needed for backtracking, making it unnecessary to restore the path of the alignment. To lower the memory usage from O($M \times N$), we split the algorithm into four main steps. In the preliminary step, we precomputed the hybridization energy for all possible pairs of ideally complementary $k$-mers based on the table of stacking energies obtained from (*99*). As a result, we obtain a $4^k \times 4^k$ matrix, whose elements are the free energies of the interaction of each $k$-mer with its ideal reverse complement. The pairs of $k$-mers that are not ideally complementary are given $+\inf$ energy; however, we also allow a small number of GT base pairs by listing all possible pairs of a given $k$-mer with the number of GT base pairs not greater than the given threshold (typically, two). The pre-computation of pairing energies for $k$-mers is performed only once and can be used several times afterwards. Given two input sequences, the algorithm extracts $k$-mers by and creates an index table. In the second step, PrePH scans for possible interaction sites. It looks up the matrix of precomputed $k$-mer stacking energies for each $k$-mer index in the first sequence and each $k$-mer index in the second sequence. All entries that were ideally complimentary on the first step are stored in $D$, $B$, and $S$ matrices. While all $k$-mers that are not complementary and therefore should not start an alignment are set to $+\inf$. On the third step, which implements a modified local alignment algorithm, we iterate through the sequences and elongate the stacking region in 9 different directions: one more pair of stacking nts, loop11, bulge10, bulge01, bulge20, bulge02, loop21, loop12, loop22 if we find non-zero energy value, i.e.

$$E' = D[i][j] + \min \begin{cases} stackingE[i-1][j+1][j][i] \\ bulge1E + stackingE[i-1][j+2][i-1][j+1][j][i] + kSE[J-1-k][I+k] \\ bulge1E + stackingE[i-2][j+1][j][i] + kSE[J-k][I+1+k] \\ bulge2E + TermAU([i-1][j+3],[j][i]) + kSE[J-2-k][I+k] \\ bulge2E + TermAU([i-3][j+1],[j][i]) + kSE[J-k][I+2+k] \\ loop11E[i-2][j+2][j][i][i-1][j+1] + kSE[J-1-k][I+1+k] \\ loop12E[i-2][j+3][j][i][i-1][j+1][j+2] + kSE[J-2-k][I+1+k] \\ loop12E[j][i][i-3][j+2][j+1][i-2][i-1] + kSE[J-1-k][I+2+k] \\ loop22E[i-3][j+3][j][i][i-2][i-1][j+1][j+2] + kSE[J-2-k][I+2+k] \end{cases},$$

where $E'$ is the energy to be calculated; $kSE$ is the precomputed $k$-mer stacking energy matrix; $stackingE$ is the matrix of stacking energies; $bulge1E$ is the energy of a 1-nt bulge; $bulge2E$ is energy of a 2-nt bulge; $TermAU$ is a function that penalizes for AU (or GU) terminating helix; $loop11E$ is the matrix of 1×1 loop energies; $loop12E$ is the matrix of 1×2 loops energies; $loop21E$ is the matrix of 2×1 loop energies; $loop22E$ is the matrix of 2×2 loop energies; $I = M - k - i$; $J = N - j - 1$.

The calculation of the energy is based on the ViennaRNA package algorithm for interior loops (*99*) with some modifications. The energy parameters are derived from the Turner energy model (*122*). Stem energies are calculated as the base pairs stacking energy of two consecutive nucleotides. The energies of bulges and interior loops usually contain a length-dependent term. Additionally, we allow for a loop structure or a bulge structure only if $k$ preceding and succeeding nucleotides are complementary. This is implemented as an additional term in the equation that extracts $k$-mer stacking energy from the pre-computed matrix. The algorithm compares the obtained new energy with global minimal energy and updates it if necessary. It also stores the start and the end coordinates of the most energy-efficient path.

In this implementation, PrePH needs only marginally more time to identify suboptimal interactions on the fourth step. We limit our search to the suboptimal structures that don't intersect. Two structures are called non-intersecting if the complementary regions of both sequences of one structure do not have nucleotides in common with complementary regions of both sequences of the second structure (it is still allowed to have intersecting regions in only one of the two sequences). To obtain the suboptimal structures, the program sets to zeros the rectangle corresponding to the optimal alignment that was already found, looks for new minimal energy in the $D_{ij}$ matrix, and checks whether the corresponding alignment intersects the optimal one. If it does, the energies along this path are set to zero. If not, the alignment is returned. These steps are repeated until all the energies below the energy threshold are considered.

The program returns a table of unique identifiers for every predicted RNA structure, its coordinates in the genome, energy in kcal/mol, and structure in dot-bracket notation. The whole pipeline is implemented as a series of python scripts and allows parallel implementation.

#### Filteration of PCCRs

In order to exclude low complexity genomic regions and conserved elements with a function other than related to RNA structure, the predicted complementary regions were intersected with the following genomic regions by bedtools intersect command (*97*). The structures overlapping the intervals from the UCSC Genome Browser tRNA track (*123–127*), sno/miRNA track (*128–134*), TFBS Conserved track, which consists of Conserved Transcription Factor Binding Sites, generated using the Transfac Matrix and Factor databases (*135*), and RepeatMasker track (*136*) (the categories “Simple repeat”, “rRNA”, “Low complexity”, “snRNA”, “srpRNA”, “DNA”) were removed. Additionally, we confined our search to RNA structures within protein-coding transcripts of protein-coding genes.

### Supplementary Figures

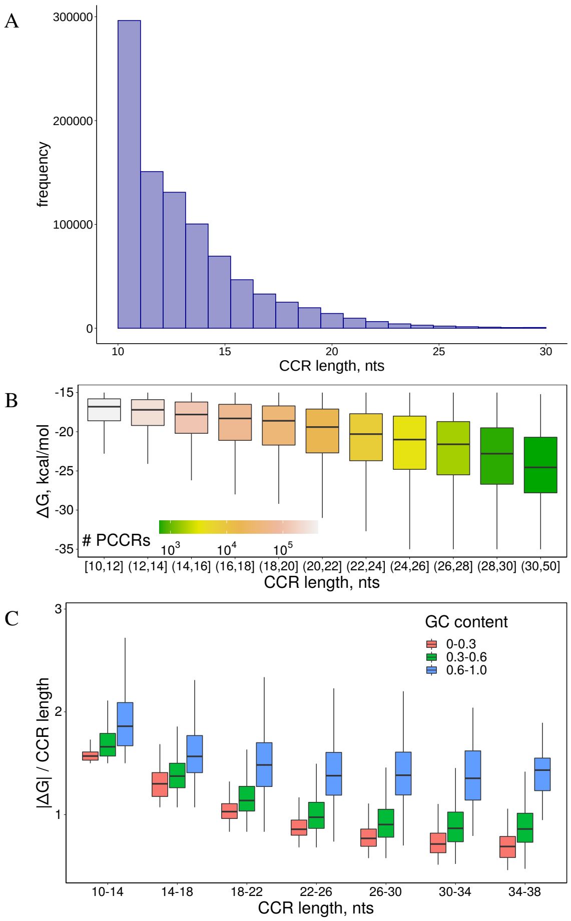

Figure S1: PCCR energy and length. **(A)** The distribution of CCR lengths. **(B)** The distribution of PCCR energies in different length groups. Colors represents the number of PCCRs in each length group. **(C)** The density of PCCR energy ($\Delta G$ per CCR length) as a function of GC content.

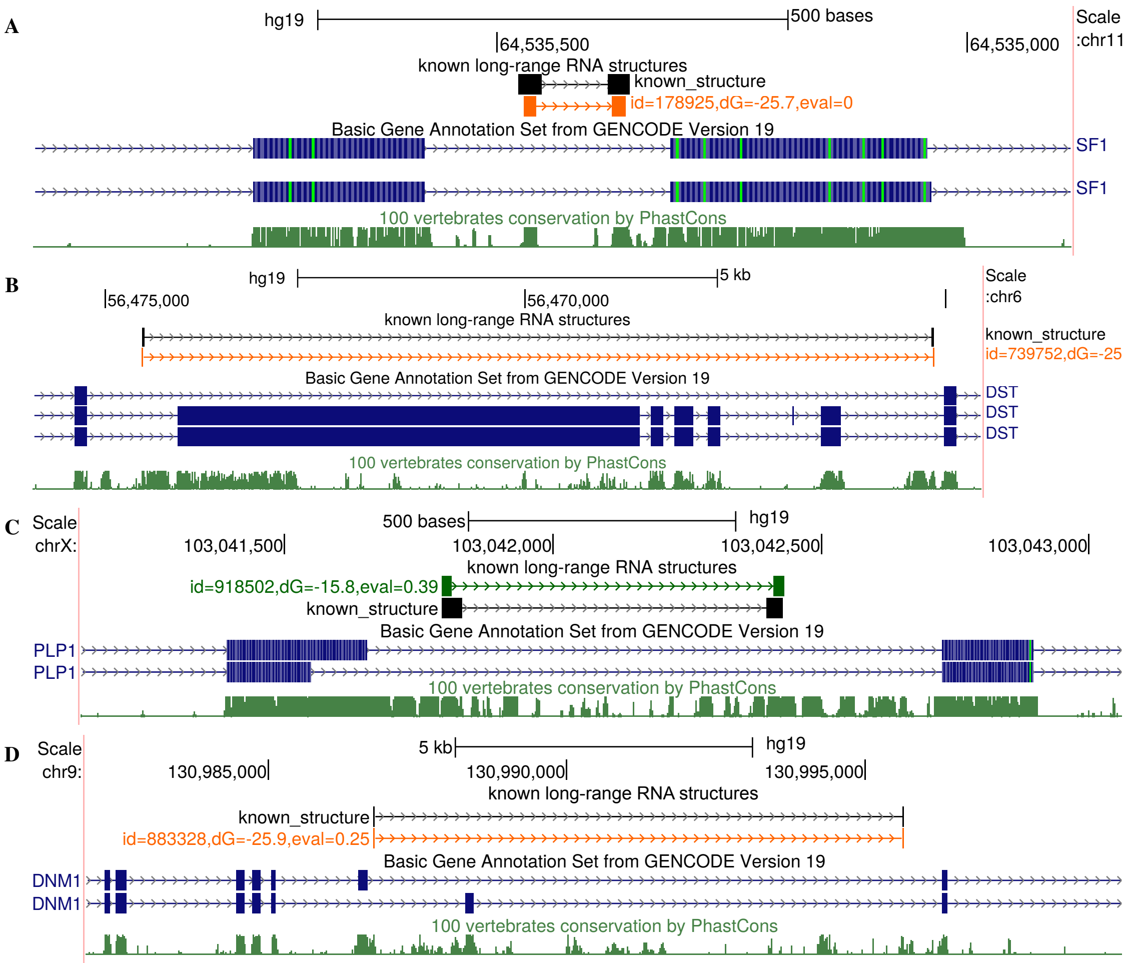

Figure S2: Long-range RNA structures in *SF1*, *DST*, *PLP1*, and *DNM1* from earlier reports (Table 1). The structure in *ENAH* gene is shown separately in Figure 6A

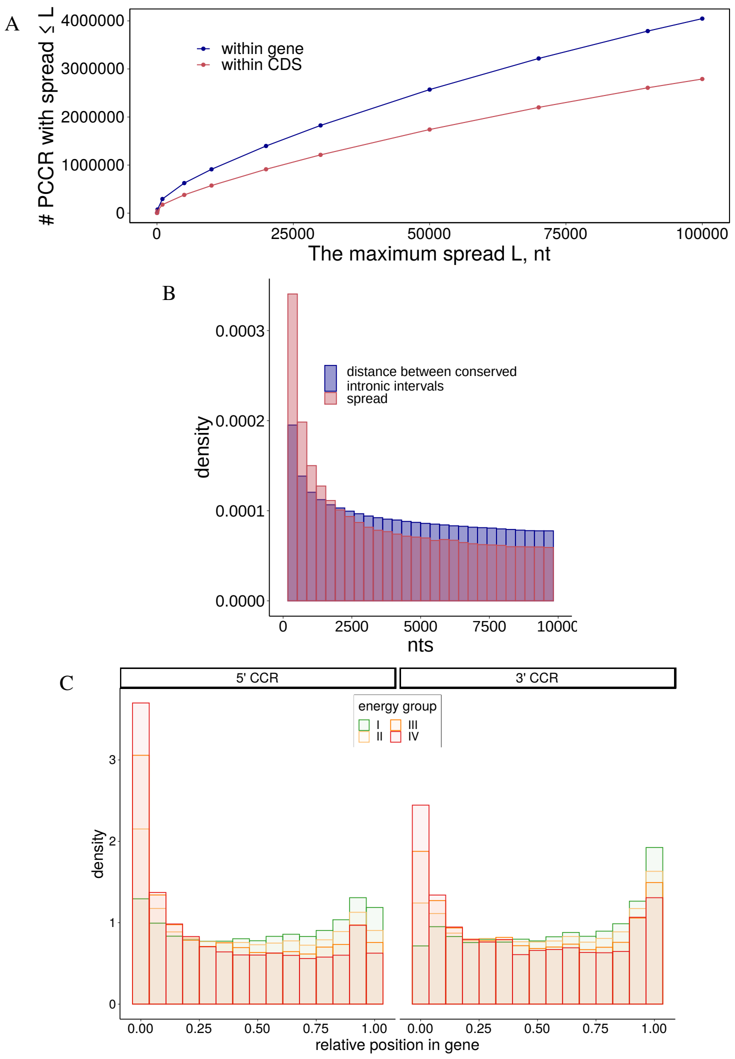

Figure S3: Properties of PCCRs. **(A)** The number of PCCRs as a function of maximum distance between CIR ($L$) in whole genes and within protein-coding parts. **(B)** The distribution of spreads and the distribution of pairwise distances between CIR. **(C)** The distribution of CCR positions along the gene.

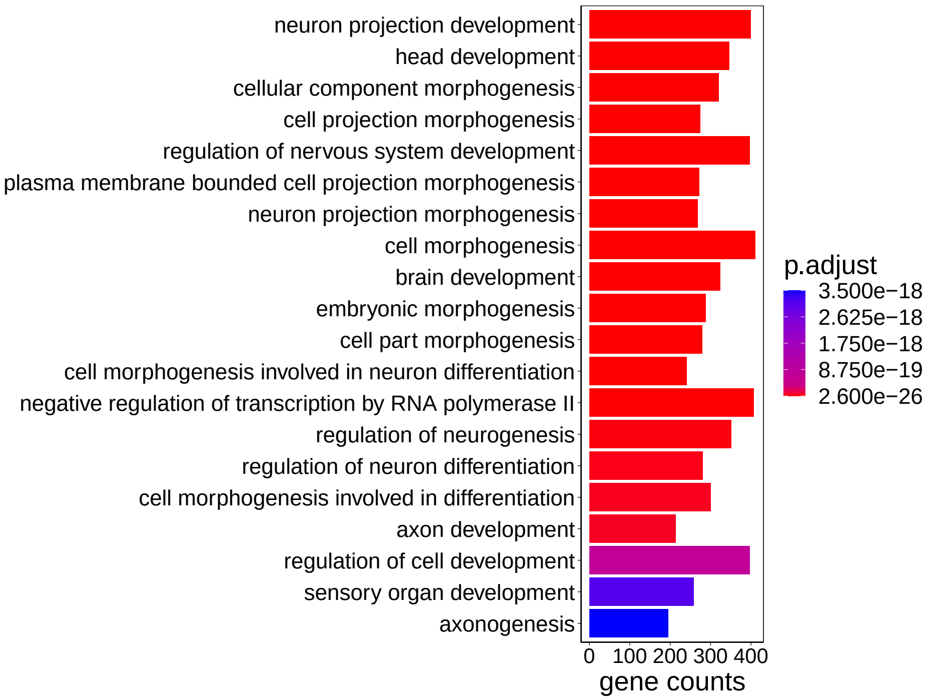

Figure S4: Enrichment of GO terms in genes with PCCRs as compared to genes matched by length, but without PCCRs.

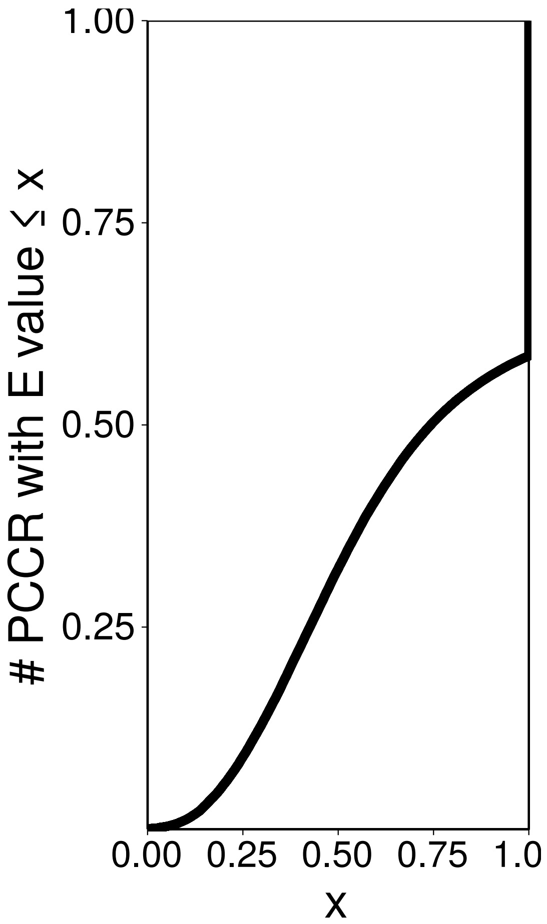

Figure S5: Cumulative distribution of E-values.

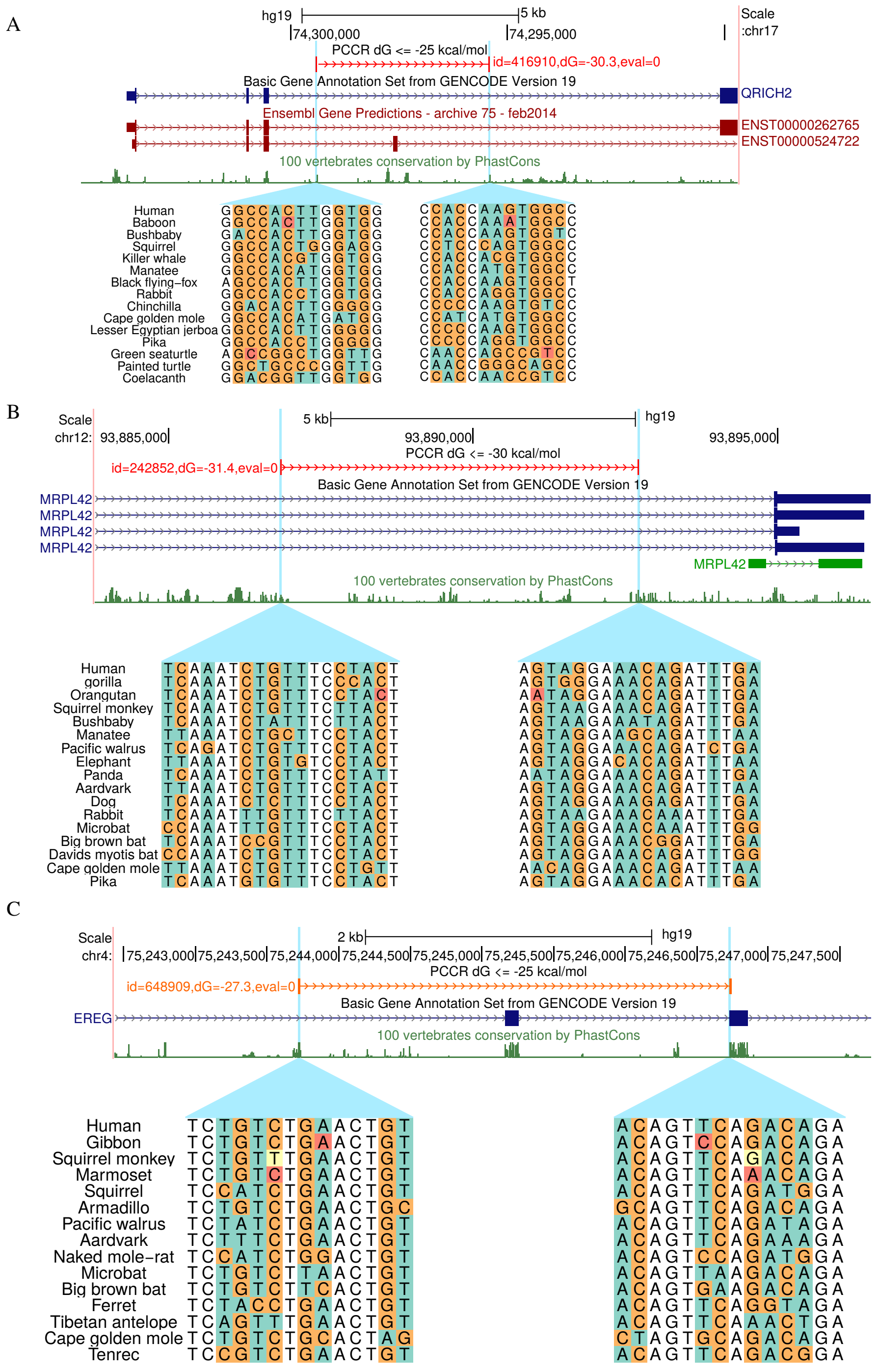

Figure S6: Structural alignments for PCCRs in the Glutamine-rich protein 2 gene (A), the 28S ribosomal protein L42 gene (B), and Epiregulin (C).

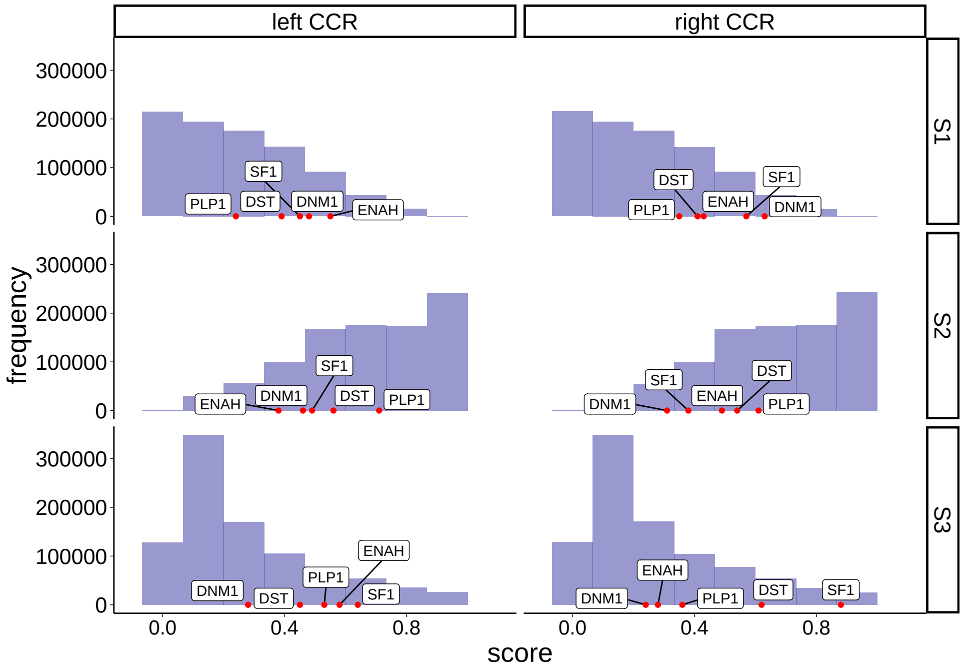

Figure S7: Conservation-derived scores. $s_1$ is the difference between the average phastCons scores within CCR and within 300 nt flanking regions (the larger, the more significant). $s_2$ is the average phastCons score within CCR and 300 nt around it (the smaller, the more significant). $s_3$ is the length of a CCR relative to the length of its parent CIR (the larger, the more significant). The respective values for *bona fide* RNA structures are shown by red dots.

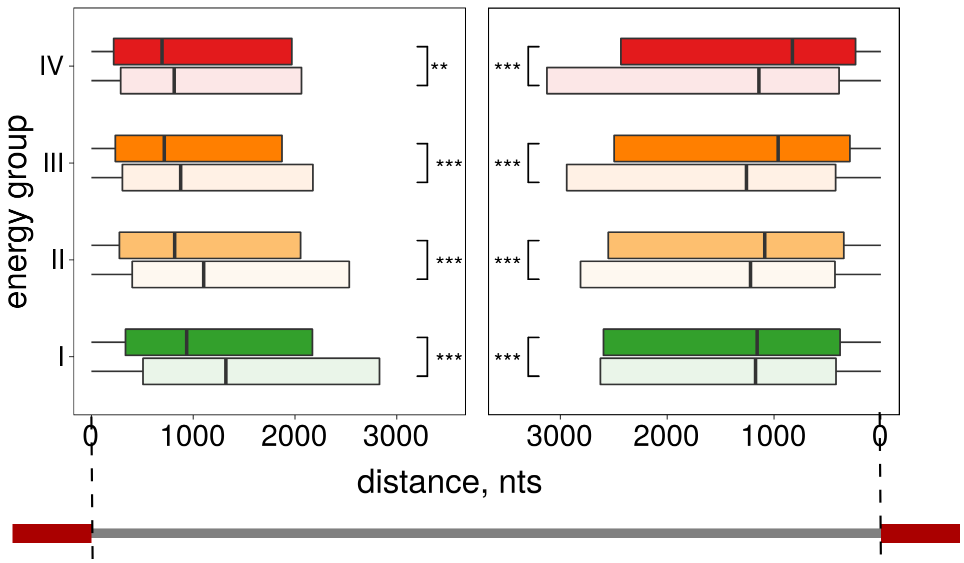

Figure S8: CCRs are located on average closer to intron ends than intervals obtained by random shift. Dark colors correspond to energy groups in Figure 3. Bright colors correspond to the respective random shift controls.

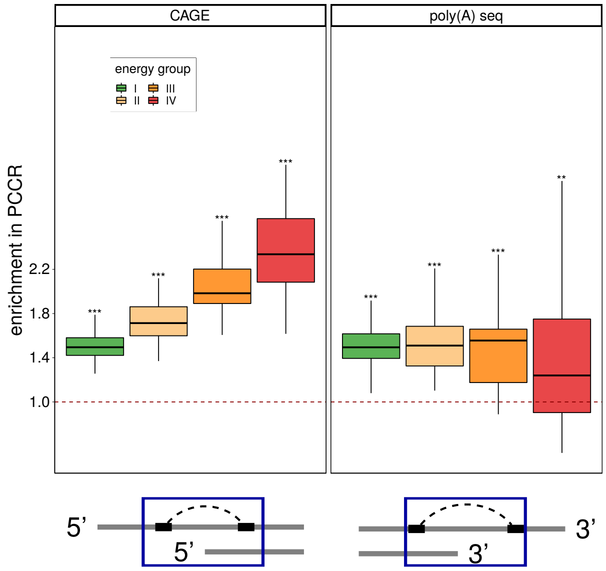

Figure S9: Expressed CAGE-tags and poly(A)-seq clusters are enriched within PCCRs.

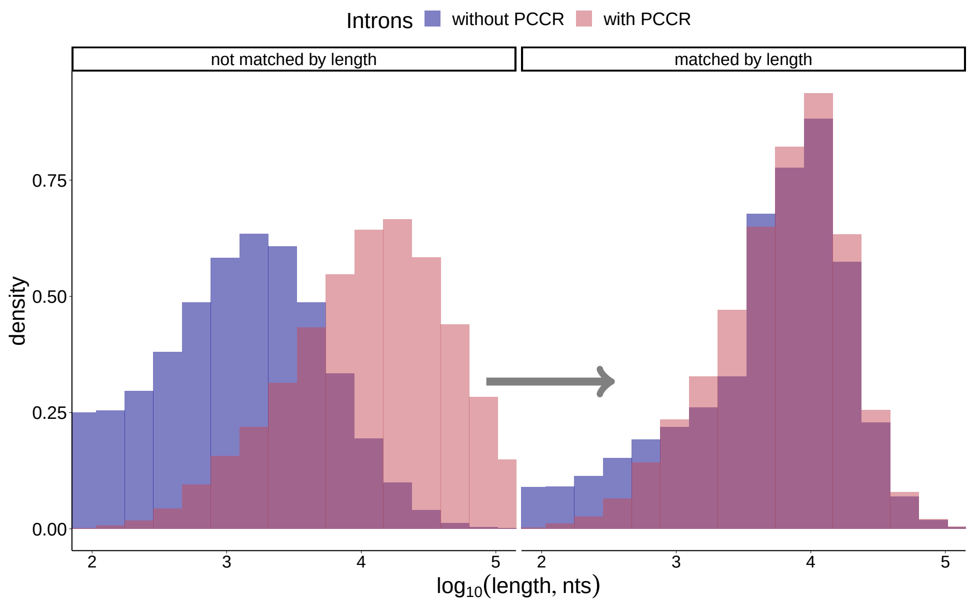

Figure S10: Matched Intron Lengths

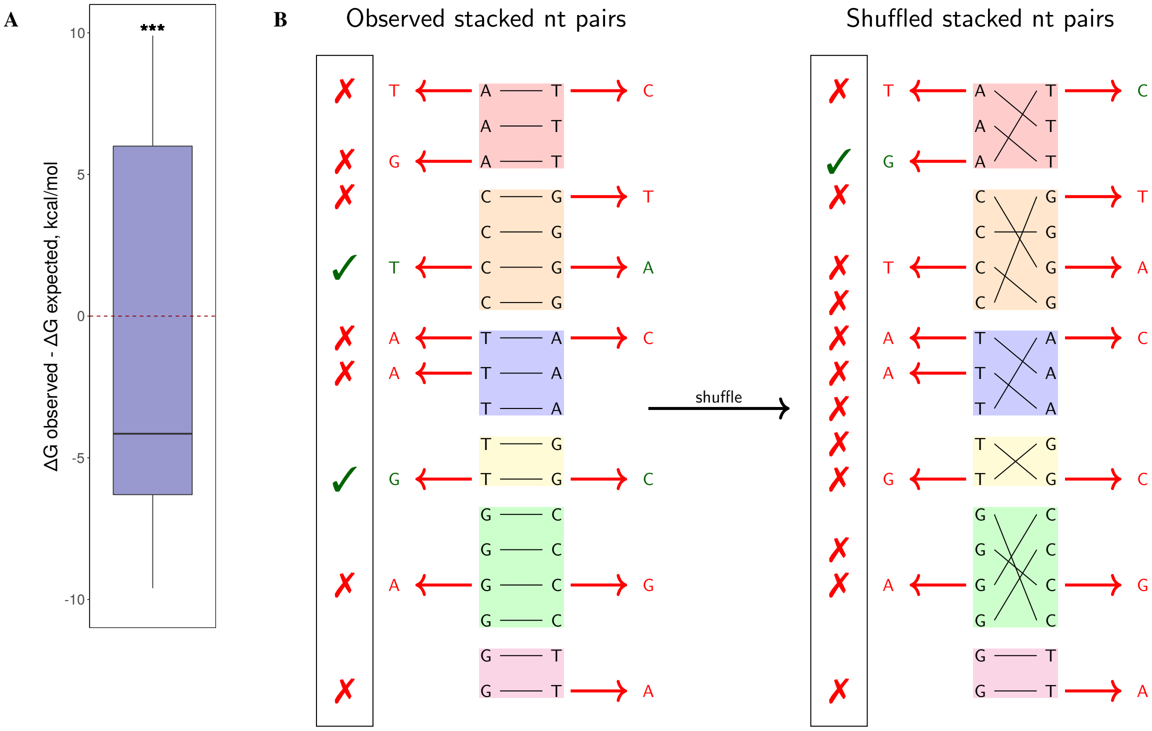

Figure S11: Mutations. (**A**) Change of $\Delta G$ in PCCRs with compensatory mutations. (**B**) Estimation of enrichment with compensatory mutations.

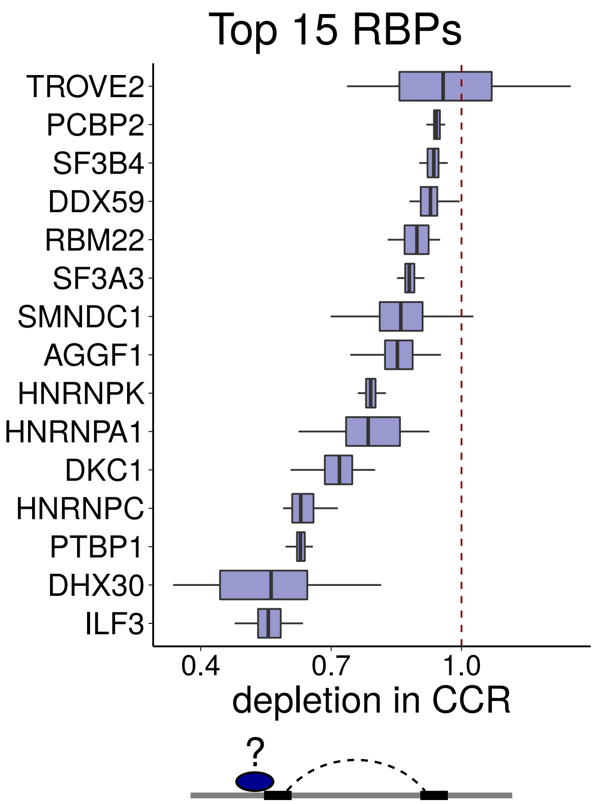

Figure S12: Top 15 RBPs with the most depleted binding within CCRs.

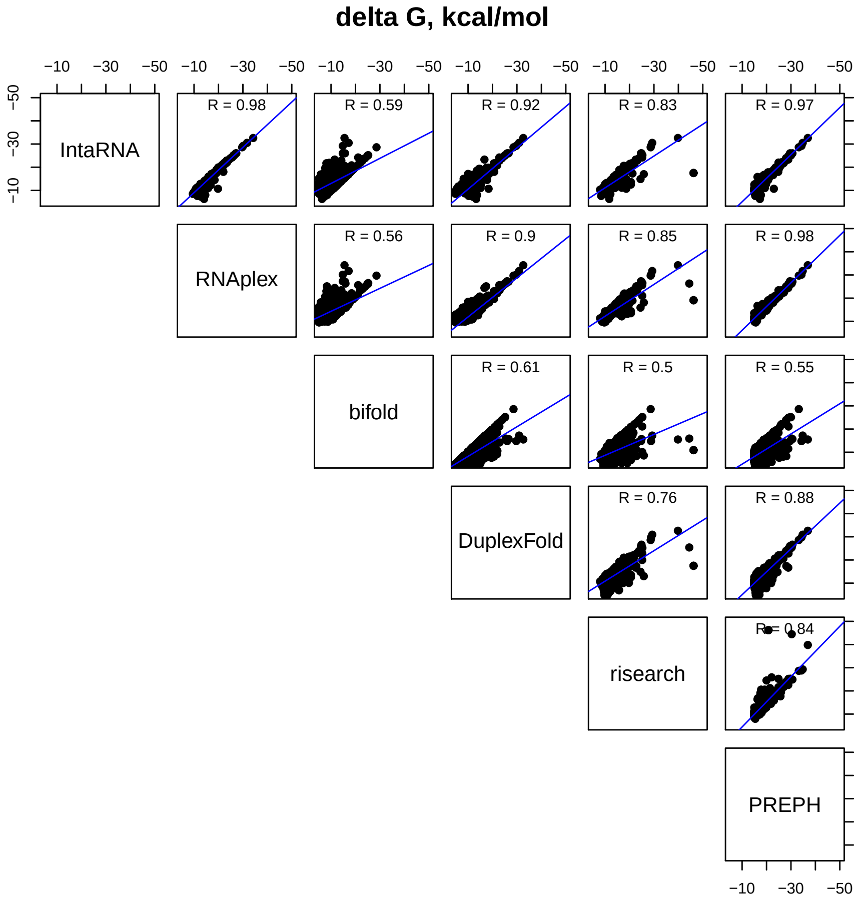

Figure S13: Correlation of predicted minimum free energies (MFE).

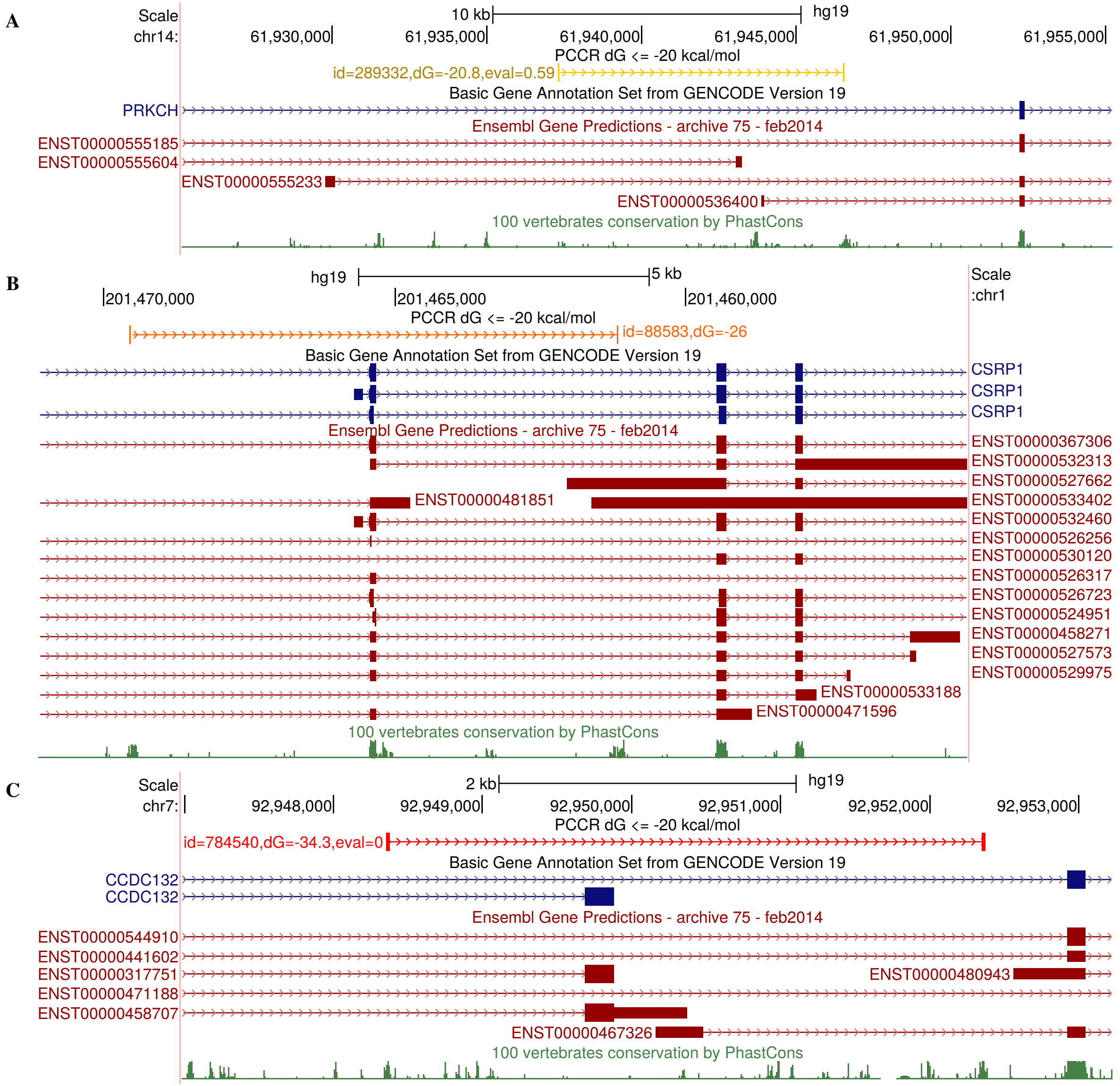

Figure S14: Transcript starts and ends inside PCCRs.

#### Supplementary Tables

| Gene | Location | Approx. coordinates | Reason not found | References |
|---|---|---|---|---|
| SMN2, SMN1 | exon 7 | chr5:70,247,788-70,248,263 | only 8-nt-long | (*137–140*) |
| TERT | exons 7 and 8 | chr5:1,271,103-1,277,432 | CCR in repeats | (*141*) |
| NEAT1 | hNEAT1-S | chr11:65,168,509-65,233,788 | lncRNA | (*142*) |
| MAPT (tau) | exon 10 | chr17:44,087,754-44,087,801 | only 6-nt-long | (*143–146*) |
| FGFR2 | stem after exon IIIb | chr10:123,277,117-123,278,208 | $\Delta G = -14.6$ kcal/mol | (*147*) |
| TNNT2 (cTNT) | exon 5 | chr1:201,338,988-201,339,019 | only 8-nt-long and no k=5 | (*148*) |
| FTL | iron RE | chr19:49,468,567-49,468,639 | outside CIR | (*149*) |
| CFTR | exon 11 | chr7:117,199,585-117,199,685 | lies in exon | (*150*) |
| GH1 (HGH) | stem before exon 3 | chr17:61,995,469-61,995,577 | multiple bulges in a row | (*78*) |
| PSEN2 | stem-loop in exon 5 | chr1:227,073,272-227,073,294 | lies in exon, too short | (*151*) |
| FN1 | stem-loop in EDA exon | chr2:216,245,605-216,245,671 | lies in exon, too short | (*152*) |
| COL2A1 | stem-loop following a weak 5'-ss | chr12:48,393,652-48,393,708 | $\Delta G = -13.9$ kcal/mol | (*153*) |
| ATE1 | exon 7b | chr10:123,631,345-123,659,239 | $d = 27,856 > 10,000$ | (*36*) |

Table S1: Experimentally-validated (*bona fide*) RNA structures in human genes that didn't satisfy search criteria. Gene name, location within the gene, and approximate coordinates in GRCh37 human genome assembly are listed.

| PREPH | RNAplex | DuplexFold | risearch | IntaRNA | bifold |
|---|---|---|---|---|---|
| 191.4 | 218.2 | 234.2 | 279.6 | 449.5 | 1301.2 |

(A)

| | PREPH | IntaRNA | RNAplex | risearch | DuplexFold | bifold |
|---|---|---|---|---|---|---|
| PREPH | 1 | 0.81 | 0.85 | 0.8 | 0.84 | 0.61 |
| IntaRNA | 0.67 | 1 | 0.86 | 0.76 | 0.72 | 0.56 |
| RNAplex | 0.72 | 0.86 | 1 | 0.8 | 0.77 | 0.5 |
| risearch | 0.68 | 0.76 | 0.8 | 1 | 0.67 | 0.48 |
| DuplexFold | 0.79 | 0.8 | 0.85 | 0.74 | 1 | 0.57 |
| bifold | 0.78 | 0.83 | 0.78 | 0.73 | 0.77 | 1 |

(B)

Table S2: Time and accuracy of PrePH with respect to IntaRNA2.0, RIsearch2, RNAplex, DuplexFold, and bifold. **(A)** The run time (sec) on 1000 pairs of randomly chosen conserved intronic sequences from the human genome. **(B)** The number of base pairs that were common between PrePH and each of the other programs as a fraction of the number of base pairs predicted by PrePH alone.

| Target | KD1 | KD2 | Ctrl1 | Ctrl2 |
|---|---|---|---|---|
| AARS | ENCFF359DJN | ENCFF809IPD | ENCFF572NPL | ENCFF689YCR |
| AATF | ENCFF283ZEU | ENCFF510KSY | ENCFF057JNE | ENCFF294SKR |
| ABCF1 | ENCFF512RCL | ENCFF375JLH | ENCFF236XCF | ENCFF901WHO |
| ACO1 | ENCFF493XEK | ENCFF974YWT | ENCFF566XUG | ENCFF486GXG |
| ADAR | ENCFF437CZI | ENCFF827TVS | ENCFF784SML | ENCFF487UNM |
| AGO1 | ENCFF888QAR | ENCFF018TKU | ENCFF057JNE | ENCFF294SKR |
| AKAP1 | ENCFF446GNQ | ENCFF554EIV | ENCFF989TXQ | ENCFF558ZFO |
| AKAP8 | ENCFF718DSO | ENCFF762NBR | ENCFF093UUE | ENCFF870WUB |
| AKAP8L | ENCFF963ZJZ | ENCFF859HYF | ENCFF566XUG | ENCFF486GXG |
| APOBEC3C | ENCFF833UXW | ENCFF360ASR | ENCFF869IKC | ENCFF996UTY |
| ASCC1 | ENCFF037MOS | ENCFF948YWB | ENCFF989TXQ | ENCFF558ZFO |
| ATP5C1 | ENCFF924MNW | ENCFF120KHG | ENCFF236XCF | ENCFF901WHO |
| AUH | ENCFF447RFE | ENCFF326XLG | ENCFF742NXL | ENCFF340UAV |
| BCCIP | ENCFF085YWZ | ENCFF813CJQ | ENCFF572NPL | ENCFF689YCR |
| BCLAF1 | ENCFF878AAZ | ENCFF587FCY | ENCFF572NPL | ENCFF689YCR |
| BOP1 | ENCFF380PZQ | ENCFF570FGE | ENCFF738YFT | ENCFF293RIK |
| BUD13 | ENCFF944UDR | ENCFF526PAV | ENCFF236XCF | ENCFF901WHO |
| CALR | ENCFF564GBC | ENCFF862YLD | ENCFF433MPP | ENCFF650KXR |
| CCAR1 | ENCFF874TAD | ENCFF981UXB | ENCFF930OYV | ENCFF934UWJ |
| CCAR2 | ENCFF867UDD | ENCFF734UFY | ENCFF261BLR | ENCFF966QBV |
| CEBPZ | ENCFF347KWA | ENCFF022CJU | ENCFF236XCF | ENCFF901WHO |
| CELF1 | ENCFF529SHX | ENCFF965ZBQ | ENCFF566IXO | ENCFF394YXV |
| CIRBP | ENCFF719TWR | ENCFF270FIC | ENCFF989TXQ | ENCFF558ZFO |
| CKAP4 | ENCFF348ABY | ENCFF855TVS | ENCFF566XUG | ENCFF486GXG |
| CNOT7 | ENCFF784QPD | ENCFF711NKA | ENCFF869IKC | ENCFF996UTY |
| CPSF6 | ENCFF183DNU | ENCFF657FCV | ENCFF738YFT | ENCFF293RIK |
| CPSF7 | ENCFF221ICX | ENCFF359NWF | ENCFF236XCF | ENCFF901WHO |
| CSTF2 | ENCFF938DQM | ENCFF866AHO | ENCFF572NPL | ENCFF689YCR |
| CSTF2T | ENCFF334UJK | ENCFF529NFO | ENCFF057JNE | ENCFF294SKR |
| DAZAP1 | ENCFF535DJP | ENCFF427XIT | ENCFF566IXO | ENCFF394YXV |
| DDX1 | ENCFF727IBQ | ENCFF902WUX | ENCFF784SML | ENCFF487UNM |
| DDX19B | ENCFF022GNJ | ENCFF430RBN | ENCFF566XUG | ENCFF486GXG |
| DDX21 | ENCFF026VLT | ENCFF332RXE | ENCFF057JNE | ENCFF294SKR |

*Continued on next page*

Table S3 – *Continued from previous page*

| Target | KD1 | KD2 | Ctrl1 | Ctrl2 |
|---|---|---|---|---|
| DDX24 | ENCFF327HAD | ENCFF805KIS | ENCFF057JNE | ENCFF294SKR |
| DDX27 | ENCFF414WJD | ENCFF210PWB | ENCFF572NPL | ENCFF689YCR |
| DDX28 | ENCFF187ZFL | ENCFF875KOM | ENCFF572NPL | ENCFF689YCR |
| DDX3X | ENCFF572OTL | ENCFF196CGJ | ENCFF989TXQ | ENCFF558ZFO |
| DDX47 | ENCFF487JHR | ENCFF161KIE | ENCFF357IOR | ENCFF239JNY |
| DDX5 | ENCFF980OCO | ENCFF435DYK | ENCFF045COW | ENCFF668JLY |
| DDX52 | ENCFF608NGH | ENCFF653INZ | ENCFF574DLW | ENCFF847SRU |
| DDX55 | ENCFF876ECR | ENCFF193SMU | ENCFF572NPL | ENCFF689YCR |
| DDX59 | ENCFF228NAK | ENCFF802SDT | ENCFF738YFT | ENCFF293RIK |
| DDX6 | ENCFF365JNK | ENCFF725RWM | ENCFF989TXQ | ENCFF558ZFO |
| DHX30 | ENCFF751TAL | ENCFF402UGA | ENCFF784SML | ENCFF487UNM |
| DKC1 | ENCFF572VRD | ENCFF454QYL | ENCFF566XUG | ENCFF486GXG |
| DNAJC2 | ENCFF534TBB | ENCFF878DOO | ENCFF784SML | ENCFF487UNM |
| DNAJC21 | ENCFF998JKT | ENCFF168QYA | ENCFF784SML | ENCFF487UNM |
| EEF2 | ENCFF832THG | ENCFF477MYS | ENCFF574DLW | ENCFF847SRU |
| EFTUD2 | ENCFF039BLK | ENCFF149WKD | ENCFF574DLW | ENCFF847SRU |
| EIF2S1 | ENCFF064LCJ | ENCFF776MXG | ENCFF989TXQ | ENCFF558ZFO |
| EIF2S2 | ENCFF079OVB | ENCFF642MGZ | ENCFF236XCF | ENCFF901WHO |
| EIF3D | ENCFF713UFE | ENCFF708DON | ENCFF574DLW | ENCFF847SRU |
| EIF3G | ENCFF794TBV | ENCFF808FDG | ENCFF057JNE | ENCFF294SKR |
| EIF4A3 | ENCFF588VRV | ENCFF160XLT | ENCFF989TXQ | ENCFF558ZFO |
| EIF4B | ENCFF400ARN | ENCFF002YLP | ENCFF989TXQ | ENCFF558ZFO |
| EIF4G1 | ENCFF756IDB | ENCFF563CZX | ENCFF566IXO | ENCFF394YXV |
| EIF4G2 | ENCFF131UOO | ENCFF028ZEH | ENCFF566IXO | ENCFF394YXV |
| ESF1 | ENCFF919BHG | ENCFF843LUG | ENCFF236XCF | ENCFF901WHO |
| ETF1 | ENCFF700QOG | ENCFF086UVA | ENCFF742NXL | ENCFF340UAV |
| EWSR1 | ENCFF398AQF | ENCFF084OLA | ENCFF566XUG | ENCFF486GXG |
| EXOSC9 | ENCFF793ZQF | ENCFF613CSR | ENCFF057JNE | ENCFF294SKR |
| FAM120A | ENCFF784QKI | ENCFF143FVG | ENCFF574DLW | ENCFF847SRU |
| FASTKD1 | ENCFF393OPX | ENCFF648JHO | ENCFF869IKC | ENCFF996UTY |
| FASTKD2 | ENCFF850TNI | ENCFF330THZ | ENCFF057JNE | ENCFF294SKR |
| FIP1L1 | ENCFF127EMC | ENCFF253SZA | ENCFF236XCF | ENCFF901WHO |
| FKBP4 | ENCFF982ALW | ENCFF257YRE | ENCFF572NPL | ENCFF689YCR |

*Continued on next page*

Table S3 – *Continued from previous page*

| Target | KD1 | KD2 | Ctrl1 | Ctrl2 |
|---|---|---|---|---|
| FMR1 | ENCFF116MWT | ENCFF913XXV | ENCFF566IXO | ENCFF394YXV |
| FTO | ENCFF408FIB | ENCFF098ONQ | ENCFF738YFT | ENCFF293RIK |
| FUBP3 | ENCFF513ADH | ENCFF880UOR | ENCFF926DGG | ENCFF718LIT |
| FXR1 | ENCFF445CHH | ENCFF969RLT | ENCFF989TXQ | ENCFF558ZFO |
| G3BP1 | ENCFF844ZHZ | ENCFF529THE | ENCFF989TXQ | ENCFF558ZFO |
| G3BP2 | ENCFF478UOM | ENCFF739NFG | ENCFF989TXQ | ENCFF558ZFO |
| GEMIN5 | ENCFF068KMA | ENCFF413TAT | ENCFF574DLW | ENCFF847SRU |
| GNB2L1 | ENCFF070ISC | ENCFF506XFA | ENCFF261BLR | ENCFF966QBV |
| GPKOW | ENCFF861JDA | ENCFF603KNO | ENCFF738YFT | ENCFF293RIK |
| GRSF1 | ENCFF016WDI | ENCFF506AFF | ENCFF869IKC | ENCFF996UTY |
| GRWD1 | ENCFF420WYN | ENCFF181DMT | ENCFF261BLR | ENCFF966QBV |
| HDGF | ENCFF745GYC | ENCFF456MAS | ENCFF566XUG | ENCFF486GXG |
| HNRNPA0 | ENCFF423AHV | ENCFF721QNS | ENCFF869IKC | ENCFF996UTY |
| HNRNPA1 | ENCFF350IPK | ENCFF045OGG | ENCFF784SML | ENCFF487UNM |
| HNRNPA2B1 | ENCFF606DES | ENCFF997UFK | ENCFF566IXO | ENCFF394YXV |
| HNRNPAB | ENCFF840CJZ | ENCFF651DBC | ENCFF869IKC | ENCFF996UTY |
| HNRNPC | ENCFF181LSI | ENCFF497PPO | ENCFF784SML | ENCFF487UNM |
| HNRNPD | ENCFF085OET | ENCFF830SPJ | ENCFF869IKC | ENCFF996UTY |
| HNRNPF | ENCFF438XHW | ENCFF593FFJ | ENCFF784SML | ENCFF487UNM |
| HNRNPK | ENCFF899ZBQ | ENCFF433BRP | ENCFF989TXQ | ENCFF558ZFO |
| HNRNPL | ENCFF536UGZ | ENCFF869LKX | ENCFF566IXO | ENCFF394YXV |
| HNRNPLL | ENCFF966FYM | ENCFF111KOR | ENCFF738YFT | ENCFF293RIK |
| HNRNPM | ENCFF601DUB | ENCFF240HMJ | ENCFF236XCF | ENCFF901WHO |
| HNRNPU | ENCFF292RTX | ENCFF516UIQ | ENCFF784SML | ENCFF487UNM |
| HNRNPUL1 | ENCFF134YSM | ENCFF596UTY | ENCFF686VNL | ENCFF430GWF |
| HSPD1 | ENCFF359GYZ | ENCFF471QSO | ENCFF093UUE | ENCFF870WUB |
| IGF2BP2 | ENCFF817RTM | ENCFF534YDS | ENCFF989TXQ | ENCFF558ZFO |
| IGF2BP3 | ENCFF753NXB | ENCFF441NPW | ENCFF989TXQ | ENCFF558ZFO |
| ILF2 | ENCFF150WUU | ENCFF886MQA | ENCFF784SML | ENCFF487UNM |
| ILF3 | ENCFF485RWH | ENCFF073SBH | ENCFF869IKC | ENCFF996UTY |
| KHDRBS1 | ENCFF831SBV | ENCFF813BGP | ENCFF869IKC | ENCFF996UTY |
| KHSRP | ENCFF021RRG | ENCFF310LTU | ENCFF989TXQ | ENCFF558ZFO |
| KIF1C | ENCFF165QIV | ENCFF576IMD | ENCFF566XUG | ENCFF486GXG |

*Continued on next page*

Table S3 – *Continued from previous page*

| Target | KD1 | KD2 | Ctrl1 | Ctrl2 |
|---|---|---|---|---|
| KRR1 | ENCFF497UDR | ENCFF576VLK | ENCFF093UUE | ENCFF870WUB |
| LARP4 | ENCFF890GSN | ENCFF760RVL | ENCFF930OYV | ENCFF934UWJ |
| LARP7 | ENCFF870DWP | ENCFF787GCZ | ENCFF261BLR | ENCFF966QBV |
| LIN28B | ENCFF511GIN | ENCFF446IUZ | ENCFF261BLR | ENCFF966QBV |
| LSM11 | ENCFF502SMN | ENCFF939XZO | ENCFF261BLR | ENCFF966QBV |
| MAGOH | ENCFF950HSR | ENCFF045JYP | ENCFF989TXQ | ENCFF558ZFO |
| MARK2 | ENCFF294MIQ | ENCFF692GMY | ENCFF261BLR | ENCFF966QBV |
| MATR3 | ENCFF780SYO | ENCFF246QJB | ENCFF989TXQ | ENCFF558ZFO |
| METAP2 | ENCFF619KWP | ENCFF249ZQO | ENCFF574DLW | ENCFF847SRU |
| MSI2 | ENCFF306PRN | ENCFF915LIM | ENCFF236XCF | ENCFF901WHO |
| MTPAP | ENCFF699CXV | ENCFF943JPB | ENCFF738YFT | ENCFF293RIK |
| NAA15 | ENCFF182PZY | ENCFF594GKW | ENCFF236XCF | ENCFF901WHO |
| NCBP2 | ENCFF594LYM | ENCFF083LTV | ENCFF989TXQ | ENCFF558ZFO |
| NELFE | ENCFF091TXV | ENCFF401AVH | ENCFF261RLR | ENCFF834TYC |
| NIP7 | ENCFF354LYI | ENCFF774DZT | ENCFF261BLR | ENCFF966QBV |
| NKRF | ENCFF294MHK | ENCFF100KDW | ENCFF093UUE | ENCFF870WUB |
| NOL12 | ENCFF806ZKL | ENCFF466PAF | ENCFF566IXO | ENCFF394YXV |
| NONO | ENCFF966TUI | ENCFF902KNB | ENCFF572NPL | ENCFF689YCR |
| NPM1 | ENCFF373WYF | ENCFF731QCQ | ENCFF686VNL | ENCFF430GWF |
| NSUN2 | ENCFF322GMR | ENCFF269GWO | ENCFF738YFT | ENCFF293RIK |
| NUFIP2 | ENCFF306PZW | ENCFF152DFW | ENCFF686VNL | ENCFF430GWF |
| NUP35 | ENCFF066VPP | ENCFF261DTU | ENCFF021BCS | ENCFF762FFH |
| NUSAP1 | ENCFF116WUD | ENCFF410NIR | ENCFF261BLR | ENCFF966QBV |
| PA2G4 | ENCFF687LTL | ENCFF136SOB | ENCFF574DLW | ENCFF847SRU |
| PABPC1 | ENCFF927GPT | ENCFF698UKF | ENCFF869IKC | ENCFF996UTY |
| PABPC4 | ENCFF359BAT | ENCFF560RJQ | ENCFF989TXQ | ENCFF558ZFO |
| PARN | ENCFF558REQ | ENCFF038YUS | ENCFF784SML | ENCFF487UNM |
| PCBP1 | ENCFF463AMS | ENCFF174ZQQ | ENCFF261RLR | ENCFF834TYC |
| PES1 | ENCFF333LCM | ENCFF964PAH | ENCFF566XUG | ENCFF486GXG |
| PHF6 | ENCFF858EYL | ENCFF306MQU | ENCFF261BLR | ENCFF966QBV |
| PKM | ENCFF249PUC | ENCFF474EDD | ENCFF574DLW | ENCFF847SRU |
| PNPT1 | ENCFF875SZJ | ENCFF713XBL | ENCFF869IKC | ENCFF996UTY |
| PPIG | ENCFF374SOY | ENCFF585WPZ | ENCFF093UUE | ENCFF870WUB |

*Continued on next page*

Table S3 – *Continued from previous page*

| Target | KD1 | KD2 | Ctrl1 | Ctrl2 |
|---|---|---|---|---|
| PPIL4 | ENCFF069NTO | ENCFF537PSL | ENCFF784SML | ENCFF487UNM |
| PRPF8 | ENCFF727IHN | ENCFF192LSC | ENCFF572NPL | ENCFF689YCR |
| PSIP1 | ENCFF048MHS | ENCFF412QTU | ENCFF261BLR | ENCFF966QBV |
| PTBP1 | ENCFF145TKU | ENCFF855TDM | ENCFF261RLR | ENCFF834TYC |
| PUF60 | ENCFF937YLH | ENCFF064JDU | ENCFF566IXO | ENCFF394YXV |
| PUM1 | ENCFF300LBV | ENCFF261TUX | ENCFF566IXO | ENCFF394YXV |
| PUM2 | ENCFF239VRF | ENCFF464WLH | ENCFF566IXO | ENCFF394YXV |
| PUS1 | ENCFF727ATR | ENCFF317YKN | ENCFF566XUG | ENCFF486GXG |
| QKI | ENCFF354FHT | ENCFF819VYL | ENCFF261RLR | ENCFF834TYC |
| RBFOX2 | ENCFF880HKN | ENCFF511KFD | ENCFF686VNL | ENCFF430GWF |
| RBM15 | ENCFF276WQL | ENCFF057WEZ | ENCFF278EWB | ENCFF376SXK |
| RBM17 | ENCFF993AIW | ENCFF442JOE | ENCFF686VNL | ENCFF430GWF |
| RBM22 | ENCFF030OLZ | ENCFF369NEZ | ENCFF742NXL | ENCFF340UAV |
| RBM25 | ENCFF518EYV | ENCFF322NIO | ENCFF261RLR | ENCFF834TYC |
| RBM27 | ENCFF541ADT | ENCFF241FYW | ENCFF261BLR | ENCFF966QBV |
| RBM39 | ENCFF343OAG | ENCFF942AVR | ENCFF686VNL | ENCFF430GWF |
| RBM47 | ENCFF680QZV | ENCFF597HYM | ENCFF869IKC | ENCFF996UTY |
| RCC2 | ENCFF267UFO | ENCFF706HMR | ENCFF261BLR | ENCFF966QBV |
| RECQL | ENCFF609EEV | ENCFF867UWM | ENCFF261RLR | ENCFF834TYC |
| RPL23A | ENCFF916TOV | ENCFF059YGS | ENCFF261BLR | ENCFF966QBV |
| RPLP0 | ENCFF380JZX | ENCFF781BVK | ENCFF261BLR | ENCFF966QBV |
| RPS19 | ENCFF707VTW | ENCFF366TRL | ENCFF574DLW | ENCFF847SRU |
| RPS2 | ENCFF108BIO | ENCFF259TDR | ENCFF261BLR | ENCFF966QBV |
| RPS3A | ENCFF054JEN | ENCFF392VDF | ENCFF869IKC | ENCFF996UTY |
| RPS5 | ENCFF455IHZ | ENCFF583EQH | ENCFF261BLR | ENCFF966QBV |
| RRP9 | ENCFF005NNX | ENCFF778BWF | ENCFF572NPL | ENCFF689YCR |
| SAFB2 | ENCFF117ZFN | ENCFF403DFM | ENCFF784SML | ENCFF487UNM |
| SART3 | ENCFF056ATS | ENCFF254RRV | ENCFF869IKC | ENCFF996UTY |
| SBDS | ENCFF801BRP | ENCFF642WPV | ENCFF869IKC | ENCFF996UTY |
| SERBP1 | ENCFF674ZQN | ENCFF171XGJ | ENCFF574DLW | ENCFF847SRU |
| SF1 | ENCFF586WHY | ENCFF772YRQ | ENCFF686VNL | ENCFF430GWF |
| SF3A3 | ENCFF923LZN | ENCFF677KRY | ENCFF168UDP | ENCFF321ZFN |
| SF3B1 | ENCFF349YJR | ENCFF465NDY | ENCFF738YFT | ENCFF293RIK |

*Continued on next page*

Table S3 – *Continued from previous page*

| Target | KD1 | KD2 | Ctrl1 | Ctrl2 |
|---|---|---|---|---|
| SF3B4 | ENCFF301YOD | ENCFF784ZDW | ENCFF261RLR | ENCFF834TYC |
| SFPQ | ENCFF611BJW | ENCFF402HCN | ENCFF686VNL | ENCFF430GWF |
| SLBP | ENCFF636XLW | ENCFF956LAK | ENCFF566IXO | ENCFF394YXV |
| SMN1 | ENCFF489WVZ | ENCFF576QGY | ENCFF784SML | ENCFF487UNM |
| SNRNP200 | ENCFF132QOS | ENCFF809FCF | ENCFF686VNL | ENCFF430GWF |
| SNRNP70 | ENCFF186MWG | ENCFF305IBD | ENCFF738YFT | ENCFF293RIK |
| SRFBP1 | ENCFF041MUH | ENCFF535THG | ENCFF742NXL | ENCFF340UAV |
| SRP68 | ENCFF137KFS | ENCFF187GDS | ENCFF572NPL | ENCFF689YCR |
| SRSF1 | ENCFF856VSZ | ENCFF942DHD | ENCFF261RLR | ENCFF834TYC |
| SRSF3 | ENCFF754ESC | ENCFF365PLL | ENCFF738YFT | ENCFF293RIK |
| SRSF5 | ENCFF332XZD | ENCFF154XVC | ENCFF566IXO | ENCFF394YXV |
| SRSF7 | ENCFF642TXU | ENCFF222WYC | ENCFF261RLR | ENCFF834TYC |
| SRSF9 | ENCFF754SUW | ENCFF664KDY | ENCFF261RLR | ENCFF834TYC |
| SSB | ENCFF647DVY | ENCFF105XIV | ENCFF742NXL | ENCFF340UAV |
| SSRP1 | ENCFF173IMV | ENCFF814EUR | ENCFF093UUE | ENCFF870WUB |
| STAU1 | ENCFF423PGY | ENCFF666GBG | ENCFF236XCF | ENCFF901WHO |
| STIP1 | ENCFF608WDW | ENCFF504JGG | ENCFF869IKC | ENCFF996UTY |
| SUCLG1 | ENCFF704JGS | ENCFF250DLK | ENCFF566IXO | ENCFF394YXV |
| SUGP2 | ENCFF434QDG | ENCFF622PET | ENCFF093UUE | ENCFF870WUB |
| SUPT6H | ENCFF179AAM | ENCFF387ASY | ENCFF236XCF | ENCFF901WHO |
| SUPV3L1 | ENCFF082CNV | ENCFF734OAK | ENCFF572NPL | ENCFF689YCR |
| TAF15 | ENCFF777VEO | ENCFF531USK | ENCFF261RLR | ENCFF834TYC |
| TARDBP | ENCFF759ZRY | ENCFF835ZPC | ENCFF738YFT | ENCFF293RIK |
| TBRG4 | ENCFF328RQK | ENCFF496DOM | ENCFF093UUE | ENCFF870WUB |
| TFIP11 | ENCFF683CRU | ENCFF319TPH | ENCFF572NPL | ENCFF689YCR |
| TIA1 | ENCFF298LYN | ENCFF243YSW | ENCFF261RLR | ENCFF834TYC |
| TIAL1 | ENCFF593CFB | ENCFF273JPU | ENCFF784SML | ENCFF487UNM |
| TRIM56 | ENCFF359RCT | ENCFF816AOJ | ENCFF784SML | ENCFF487UNM |
| TROVE2 | ENCFF119ZCG | ENCFF115CEE | ENCFF574DLW | ENCFF847SRU |
| TUFM | ENCFF328WOU | ENCFF709PZI | ENCFF566IXO | ENCFF394YXV |
| U2AF1 | ENCFF163IXV | ENCFF172LGC | ENCFF236XCF | ENCFF901WHO |
| U2AF2 | ENCFF911EJE | ENCFF821FZB | ENCFF686VNL | ENCFF430GWF |
| UBE2L3 | ENCFF022GVT | ENCFF416BYL | ENCFF566IXO | ENCFF394YXV |

*Continued on next page*

Table S3 – *Continued from previous page*

| Target | KD1 | KD2 | Ctrl1 | Ctrl2 |
|---|---|---|---|---|
| UCHL5 | ENCFF432SEN | ENCFF300CDS | ENCFF572NPL | ENCFF689YCR |
| UPF1 | ENCFF850ORL | ENCFF819BTX | ENCFF168UDP | ENCFF321ZFN |
| UPF2 | ENCFF023KBS | ENCFF737IEW | ENCFF572NPL | ENCFF689YCR |
| UTP18 | ENCFF977IXL | ENCFF202TGW | ENCFF168UDP | ENCFF321ZFN |
| UTP3 | ENCFF131RNR | ENCFF876DBS | ENCFF168UDP | ENCFF321ZFN |
| XPO5 | ENCFF386NUM | ENCFF234HPE | ENCFF869IKC | ENCFF996UTY |
| XRN2 | ENCFF450HWR | ENCFF789QEA | ENCFF572NPL | ENCFF689YCR |
| YBX3 | ENCFF296FTT | ENCFF036XRE | ENCFF784SML | ENCFF487UNM |
| ZNF622 | ENCFF846YSQ | ENCFF776IMC | ENCFF168UDP | ENCFF321ZFN |

Table S3: The list of accession numbers (*109, 110*) of RBP shRNA-knockdown followed by RNA-seq. Two knockdown bioreplicates (KD1, KD2) and their control experiments (Ctrl1, Ctrl2).

| Target | eCLIP1 | eCLIP2 |
|---|---|---|
| AGGF1 | ENCFF388PDG | ENCFF283MCJ |
| AKAP1 | ENCFF271UXH | ENCFF807KPM |
| BCCIP | ENCFF627YDB | ENCFF698TRT |
| BCLAF1 | ENCFF691JPV | ENCFF858HBB |
| BUD13 | ENCFF364QPG | ENCFF929ITC |
| CDC40 | ENCFF890AXT | ENCFF513MLD |
| CSTF2 | ENCFF892JUQ | ENCFF370ZBJ |
| CSTF2T | ENCFF700HIR | ENCFF786JAM |
| DDX3X | ENCFF798XQM | ENCFF799ICU |
| DDX52 | ENCFF591WOA | ENCFF539JUN |
| DDX55 | ENCFF315AVG | ENCFF584QHX |
| DDX59 | ENCFF441KPC | ENCFF485MSN |
| DDX6 | ENCFF438KSN | ENCFF650BQD |
| DGCR8 | ENCFF674MHQ | ENCFF382SXP |
| DHX30 | ENCFF506NPG | ENCFF191HRQ |
| DKC1 | ENCFF074ATJ | ENCFF043XKO |
| EFTUD2 | ENCFF511ZTQ | ENCFF955RUR |

*Continued on next page*

Table S4 – *Continued from previous page*

| Target | eCLIP1 | eCLIP2 |
|---|---|---|
| EIF3D | ENCFF650RVF | ENCFF738CPK |
| EIF3H | ENCFF770FPA | ENCFF582LGR |
| FAM120A | ENCFF511SXK | ENCFF287ADK |
| FASTKD2 | ENCFF503DBN | ENCFF136QLO |
| FKBP4 | ENCFF993XGD | ENCFF585PKG |
| FTO | ENCFF655JNU | ENCFF925SGC |
| FUBP3 | ENCFF410ITK | ENCFF714TFP |
| G3BP1 | ENCFF934JFZ | ENCFF165JYX |
| GRSF1 | ENCFF733VOV | ENCFF451GQU |
| GRWD1 | ENCFF689GDA | ENCFF735YNV |
| GTF2F1 | ENCFF092IJR | ENCFF240QYG |
| HNRNPA1 | ENCFF535MAE | ENCFF604WJZ |
| HNRNPC | ENCFF401SPQ | ENCFF738FJS |
| HNRNPK | ENCFF877KBJ | ENCFF556EBT |
| HNRNPL | ENCFF581SSQ | ENCFF415HDB |
| HNRNPM | ENCFF542FRX | ENCFF040JSC |
| HNRNPU | ENCFF024FCB | ENCFF587FNW |
| HNRNPUL1 | ENCFF676GXG | ENCFF801KHR |
| IGF2BP1 | ENCFF976DBP | ENCFF486BXN |
| IGF2BP3 | ENCFF858ZLX | ENCFF175PMW |
| ILF3 | ENCFF460MRE | ENCFF535COB |
| KHSRP | ENCFF509GWT | ENCFF891IAC |
| LARP4 | ENCFF288SGB | ENCFF130RAQ |
| LARP7 | ENCFF699ZLU | ENCFF440FLF |
| LIN28B | ENCFF976KYV | ENCFF381VVY |
| LSM11 | ENCFF870HZF | ENCFF220ECM |
| MATR3 | ENCFF874BAL | ENCFF333CUY |
| NCBP2 | ENCFF047YUZ | ENCFF500HBQ |
| NIP7 | ENCFF077RVD | ENCFF527MFS |
| NKRF | ENCFF233SNH | ENCFF864UGQ |
| NOL12 | ENCFF716SCR | ENCFF515WRY |
| PCBP1 | ENCFF252BJN | ENCFF940CJE |
| PCBP2 | ENCFF074AQN | ENCFF322DXB |

*Continued on next page*

Table S4 – *Continued from previous page*

| Target | eCLIP1 | eCLIP2 |
|---|---|---|
| PPIG | ENCFF597JSM | ENCFF603DSB |
| PRPF8 | ENCFF646XFI | ENCFF156PFJ |
| PTBP1 | ENCFF154BAE | ENCFF471GJE |
| QKI | ENCFF525PZA | ENCFF459ANM |
| RBFOX2 | ENCFF639MYI | ENCFF664WCU |
| RBM15 | ENCFF017HYK | ENCFF971BBY |
| RBM22 | ENCFF353XWS | ENCFF990UNN |
| RBM5 | ENCFF980CBO | ENCFF663GBN |
| RPS3 | ENCFF843WBE | ENCFF163XOB |
| SF3A3 | ENCFF642VTG | ENCFF075FPZ |
| SF3B4 | ENCFF581RVY | ENCFF363PGR |
| SFPQ | ENCFF221WOF | ENCFF919FIY |
| SLTM | ENCFF870XZV | ENCFF892IWC |
| SMNDC1 | ENCFF230AER | ENCFF240MCZ |
| SND1 | ENCFF054ZXE | ENCFF749TYO |
| SRSF1 | ENCFF179SCM | ENCFF184TBM |
| SRSF7 | ENCFF774EDY | ENCFF913UPA |
| SRSF9 | ENCFF125UVA | ENCFF409GPI |
| SSB | ENCFF828WCR | ENCFF746RLH |
| SUB1 | ENCFF033IXH | ENCFF978MMK |
| SUGP2 | ENCFF187DUQ | ENCFF699DTX |
| SUPV3L1 | ENCFF882KLS | ENCFF089NRQ |
| TAF15 | ENCFF603JTU | ENCFF063BTH |
| TBRG4 | ENCFF472CCD | ENCFF454ZHL |
| TIA1 | ENCFF055IIM | ENCFF358SGV |
| TIAL1 | ENCFF456DAU | ENCFF473IIG |
| TRA2A | ENCFF210QRC | ENCFF955HGE |
| TROVE2 | ENCFF515SVK | ENCFF029CYV |
| U2AF1 | ENCFF078SCZ | ENCFF295GHA |
| U2AF2 | ENCFF444TCK | ENCFF523JWI |
| UCHL5 | ENCFF305LRB | ENCFF171KTG |
| UPF1 | ENCFF848JHB | ENCFF483STL |
| UTP18 | ENCFF227SZN | ENCFF235ASG |

*Continued on next page*

Table S4 – *Continued from previous page*

| Target | eCLIP1 | eCLIP2 |
|---|---|---|
| XPO5 | ENCFF756YND | ENCFF028MVM |
| XRCC6 | ENCFF225IXQ | ENCFF381LFD |
| XRN2 | ENCFF845EBP | ENCFF944KMN |
| YBX3 | ENCFF268SII | ENCFF904UKX |

Table S4: The list of accession numbers (*109, 110*) of RBP eCLIP peaks; two bioreplicates (eCLIP1 and eCLIP2).

**SupplementaryDataFile 1:** The full list of PCCRs, GRCh37 Human Genome assembly.

http://arkuda.skoltech.ru/∼dp/shared/PrePH/SupplementaryDataFile1.bed

**SupplementaryDataFile 2:** The full list of PCCRs, GRCh38 Human Genome assembly.

http://arkuda.skoltech.ru/∼dp/shared/PrePH/SupplementaryDataFile2.bed

**SupplementaryDataFile 3:** RNA bridges, GRCh37 Human Genome assembly.

http://arkuda.skoltech.ru/∼dp/shared/PrePH/SupplementaryDataFile3.tsv

**SupplementaryDataFile 4:** Exon loop-outs, GRCh37 Human Genome assembly.

http://arkuda.skoltech.ru/∼dp/shared/PrePH/SupplementaryDataFile4.tsv

**SupplementaryDataFile 5:** A stringent set of intramolecular RIC-seq RNA contacts, provided as a courtesy of Prof. Xue (*24*).

http://arkuda.skoltech.ru/∼dp/shared/PrePH/SupplementaryDataFile5.bed
